# Characterization of the *Plasmodium berghei* regulatory AAA-ATPase subunit Rpt3 as an activator of Protein Phosphatase 1: direct and indirect evidence

**DOI:** 10.1101/2023.07.20.549862

**Authors:** Claudianne Lainé, Caroline De Witte, Alain Martoriati, Amaury Farce, Inès Metatla, Ida Chiara Guerrera, Katia Cailliau, Jamal Khalife, Christine Pierrot

**Affiliations:** Univ. Lille, CNRS, Inserm, CHU Lille, Institut Pasteur de Lille, U1019 - UMR 9017 -CIIL - Center for Infection and Immunity of Lille, F-59000 Lille, France; Univ. Lille, CNRS, UMR 8576-UGSF-Unité de Glycobiologie Structurale et Fonctionnelle, F-59000 Lille, France; University of Lille, Inserm, CHU Lille, U1286 - Infinite - Institute for Translational Research in Inflammation, F-59000 Lille, France; Proteomics platform 3P5-Necker, Université Paris Descartes - Structure Fédérative de Recherche Necker, INSERM US24/CNRS UMS3633, Paris, France

**Author notes:** These authors have contributed equally to this work (Claudianne Lainé, Caroline De Witte).

**Keywords:** proteasome, 19S regulatory particle, Rpt3, Plasmodium, PP1

## Abstract

The 26S proteasome is the main proteolytic machine involved in protein degradation, thus contributing to homeostasis or stress response of eukaryotic cells. This macromolecular complex, consisting of a 20S core particle assembled with one or two 19S regulatory particles, is highly regulated by phosphorylation. Here we describe the *Plasmodium berghei* proteasome AAA-ATPase regulatory subunit Rpt3 and show that it binds to protein phosphatase 1, the major parasite phosphatase. In addition, PbRpt3 regulates the activity of the phosphatase both in vitro and in a heterologous model of *Xenopus* oocytes. Using mutagenesis approaches, we observed that the RVXF motifs of PbRpt3 are involved in this binding and activity. Further use of *Xenopus* oocyte model and mutagenesis based on the 3D model that we established revealed that the binding capacity of PbRpt3 to ATP may also contribute to its phosphatase-regulating activity. In the parasite, reverse genetic studies suggested an essential role for PbRpt3 since no viable knock-out line could be obtained. Additionally, immunoprecipitation assays followed by mass spectrometry analyses using transgenic PbRpt3-tagged parasites not only confirmed that PbRpt3 belongs to the 19S regulatory particle of the proteasome, but also revealed potential interaction with proteins already shown to play a role in the phospholipid membrane dynamics.

## Introduction

The ubiquitin-proteasome system is one of the main pathways in protein degradation. Its cellular functions range from general cellular homeostasis and stress response to the control of vital processes such as cell division and signal transduction. The proteasome is a large multi-protein enzyme complex found in eukaryotes, archaea and some bacteria of the order Actinomycetales (for review [1]). In eukaryotic cells, it is constitutively present in the cytosol, associated with the endoplasmic reticulum, and can also be present in the nucleus, depending on the cell-type, growth status, and whether the cell is encountering stimulation or stress [2–4]. As the endpoint for the ubiquitin-proteasome system, the 26S proteasome is the principal proteolytic machine responsible for protein degradation of misfolded, denatured or obsolete proteins by proteolysis. The fully assembled 26S proteasome is a macromolecular complex consisting of a 20S core particle and one or two 19S regulatory particles. The 19S is composed of two subcomplexes, the base which directly binds to the 20S, and the lid [5]. The base is formed by six AAA-ATPase subunits (Rpt1-6), and four non-ATPase subunits (Rpn1, Rpn2, Rpn10, and Rpn13). The other nine non-ATPase subunits form the lid. The role of the 19S is to recognize, to unfold the polyubiquitinated substrates and to open the channel prior to the substrate translocation into the 20S cavity. All these activities except for substrate recognition depend on ATP binding/hydrolysis by the ATPase subunits. The core 20S particle forms a cylinder which houses the proteolytic activity of the proteasome within a central chamber [6]. It is composed of two related types of subunits; α subunits, which form the outer two heptameric rings, and β subunits, which form the inner pair of heptameric rings and include the proteolytic active sites.

The 26S proteasome is a dynamic complex highly regulated by post-translational modifications (PTMs). These PTMs modulate numerous aspects of proteasome functions, including catalytic activities, assembly and turnover of the complex, interactions with associating partners, subcellular localization, and substrate preference [7]. Among these PTMs, reversible phosphorylation is one of the most frequent and best studied PTM of the proteasome (for review [8]). Thanks to large-scale MS studies, more than 400 phosphosites have been reported on human 26S proteasome and detected on each subunit. While the kinases involved in proteasome phosphorylation have been well studied and include the protein kinases PKA, PKG and CK2, little is known about the phosphatases implicated in the counterbalance dephosphorylation reactions. Treatment of the proteasome with phosphatases such as Protein phosphatase 1 (PP1) or Protein phosphatase 2A (PP2A) reduces its activity *in vitro* [9–11]. Similarly, some PP2A and calcineurin subunits have been found complexed to the 20S proteasome [9,12,13]. However, the exact role of these phosphatases in proteasome regulation is still in its infancy. The only phosphatase that has been shown to play a physiological role is UBLCP1 (Ubiquitin-like domain containing CTD phosphatase-1), which negatively regulates proteasome function by promoting 19S-20S dissociation [14].

More recently, our analyses based on a proteomic approach conducted in *Plasmodium berghei* (Pb) have identified some of the 19S AAA-ATPase and non-ATPase subunits as potential interactors of the parasite PP1 catalytic subunit (PP1c) [15,16]. This parasite, causative agent of malaria, undergoes rapid growth and cell division in an oxidatively stressed environment. Consequently, it is highly dependent of a tight and quick protein turnover machinery. Interestingly, proteasome inhibitors exhibit parasiticidal activity at different stages of the parasite’s lifecycle (for review [17]). Moreover, inhibitors of the proteasome strongly synergize artemisinin-induced killing of *Plasmodium*, both *in vitro* and *in vivo* [18,19]. In *Plasmodium falciparum* (Pf), the proteasome 26S has been purified and characterized, together with its physiologic interacting partners [20,21]. As in eukaryotes, it is composed of a 19S regulatory particle and a 20S core particle, the structure of which has been solved by high-resolution electron cryo-microscopy, thus establishing the molecular basis of the parasite 20S proteasome specificity, when compared to its human counterpart [22].

As for PP1, it has been shown to be a key enzyme in *Plasmodium*. This phosphatase seems to be responsible for most of the dephosphorylation processes in the parasite [23], and reverse genetic analyses have demonstrated its essentiality for blood-stage parasite development and egress from erythrocytes [24,25]. PP1 is a holoenzyme composed of a highly conserved catalytic subunit (PP1c) in association with one or several regulatory subunits which direct its localization and shape its activity/specificity [26–30]. A common feature of most of the PP1c regulators is that they contain a RVxF sequence as the primary PP1c-binding motif [31]. In *Plasmodium*, numerous potential regulatory partners have been identified in global analyses of PP1c interactome [15,16] . This includes both conserved and specific regulators [15,32–35]. Among the PP1c-binding proteins, some components of the proteasome 19S were identified, suggesting that they could be PP1c substrates, recruit additional substrates, transport PP1c, and/or regulate the phosphatase activity.

In this paper, we report the molecular and functional characterization of PbRpt3, one of the *P. berghei* 19S AAA-ATPase subunits. We showed that it binds to PP1c and increases the phosphatase activity both *in vitro* and in the heterologous model of *Xenopus* oocytes. Using site-directed mutagenesis, we demonstrated that the PbRpt3 RVxF motifs play a role in this interaction and function, and provided evidence that the enzymatic ATPase activity *per s*e of the protein is involved in its regulatory activity. Studies in the parasite suggest an essential role of PbRpt3 since no viable knock-out parasites could be obtained. Finally, by immunoprecipitation and mass spectrometry analysis, we identified many potential PbRpt3 interactors, including not only the 19S components of the proteasome, but also proteins already found to play a role in the phospholipid membrane dynamics. Based on these observations, our results unravel PbRpt3 as a new activator of PP1c with a function linked to its enzymatic activity, and this ATPase is likely to be involved in a newly described pathway unrelated to the 20S catalytic role of the proteasome and linked to membrane phospholipid binding.

## Results and Discussion

### PbRpt3 sequence analysis and protein annotation

Exploration of PlasmoDB database showed that PbRpt3 gene (PBANKA_0715600) is predicted to contain 1516 bp, including an intron of 328 bp. The sequencing of the PCR products obtained with specific primers on both cDNA and genomic DNA confirmed both the transcription of PbRpt3 and its correct exons-introns positions respectively (data not shown). The deduced sequence of 395 amino acid (a.a.) (Supplementary Figure 1) showed 100% identity to the PlasmoDB predicted sequence. Although the *Plasmodium* protein is 23 amino acids shorter that its human counterpart PSMC4 (Supplementary Figure 2), it exhibits the same regions and domains. Indeed, the InterproScan analysis of PbRpt3 potential domains revealed the presence of an AAA-ATPase domain with a Walker A motif (consensus G-x(4)-GK-[TS] and sequence GPPGTGKT) at position 182 and a Walker B motif (consensus hhhDE and sequence IIFIDE) at position 237 (Figure 1A, Supplementary Figure 1). These ATP binding motifs are also present in human Rpt [36]. A coil domain is predicted at the Nter of PbRpt3. Coil domains have been previously described to be involved in the coiled-coil interaction between the different Rpt proteins of the proteasome 19S complex [37]. Further, a.a. sequence analysis showed the presence of two consensus RVxF motifs, well known as PP1c binding motif. These two motifs, at positions 200 (200-KVTF-203) and 305 (305-RKIEF-309), fit to the consensus sequence [R/K]-X(0-1)- [V/I]-{P}-[F/W] ({P} being any amino acid except a proline) (Figure 1B). The first motif KVTF is not conserved between all *Plasmodium* species (Supplementary Table 1). It is present in *Plasmodium knowlesi* Rpt3 but absent in *Plasmodium falciparum* Rpt3 (PF3D7_0413600) and three other *Plasmodium* species infecting humans. Regarding the second motif RKIEF, it is conserved among all *Plasmodium* species as well as among mammals (*Homo sapiens, Mus musculus*), yeast (*Saccharomyces cerevisiae)* and plants (*Arabidopsis thaliana* - Uniprot Reference A0A178UAH2). The presence of two putative RVxF motifs in PbRpt3 may explain its detection in PbPP1c interactome [15,16], and likely suggests that it could be a direct interactor of the phosphatase.

**FIGURE 1.**
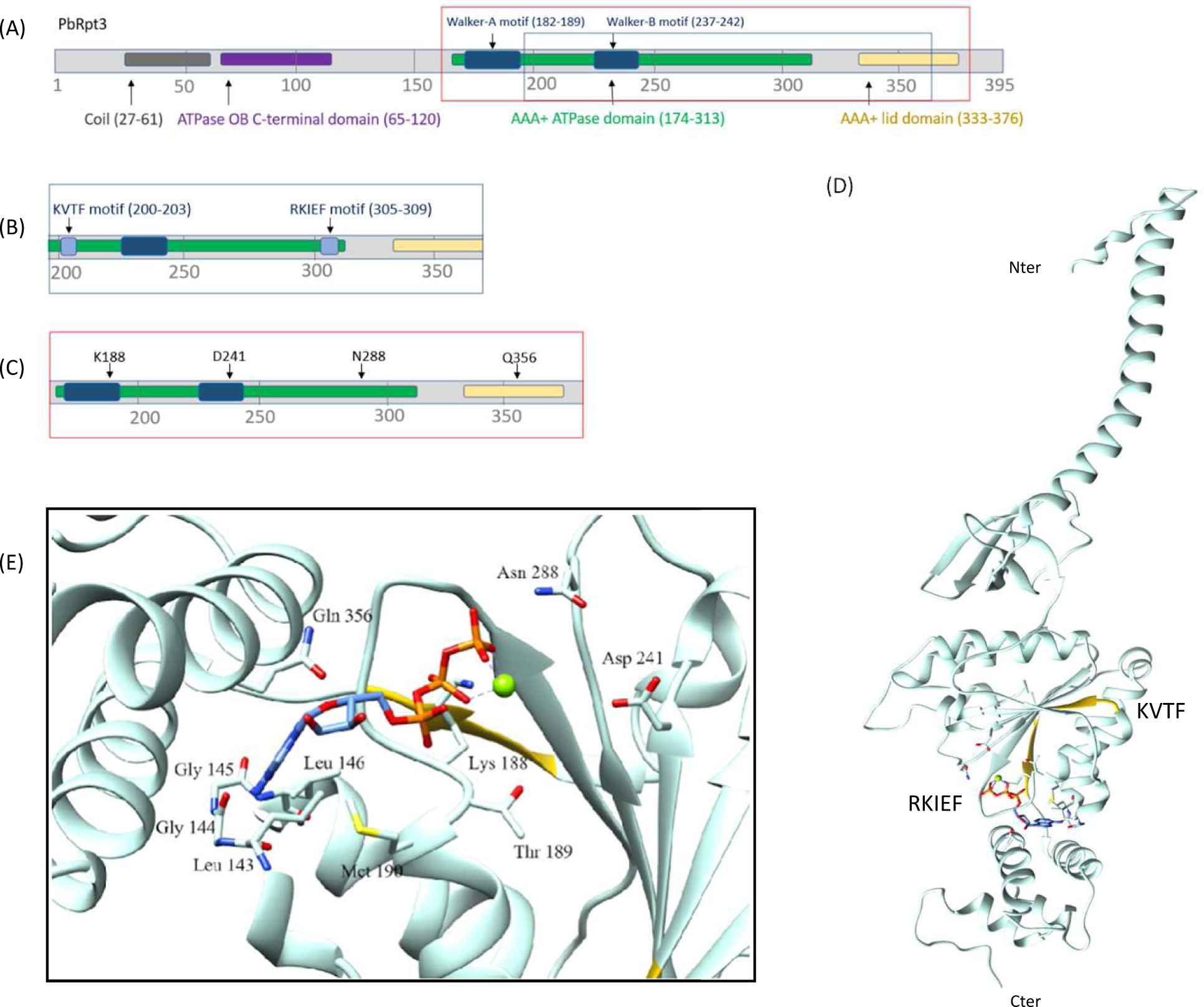
*Plasmodium berghei* regulatory AAA-ATPase subunit PbRpt3 domains and structural model. **A.** Prediction of the PbRpt3 domains using InterproScan software. Schematic of the PbRpt3 domains showing **B.** part of the AAA-ATPase domain corresponding to the two RVxF consensus sequences: KVTF and RKIEF and **C.** part of the AAA-ATPase domain where four amino acids are located: K188, D241, N288 and Q356 which are essential for Mg^2+^ and ATP binding. **D.** Predicted tertiary structure of PbRpt3. Image generated from PDB file 6MSB P auth D chain predicted with Sybyl 6.9.2 (Tripos) and annotated with Chimera, highlighting the two RVXF motifs (in yellow) and the ATP molecule at the centre. **E.** Inset shows the side structure of the amino acid chains involved in ATP binding and Mg^2+^ stabilization : the ATP molecule is modelled next to the Mg^2+^ ion (green) and surrounded by K188, and Q356 which are the main holders of the ATP molecule.

### PbRpt3 exhibits ATP-binding ability and would require a Mg2+ ion for the protein stability

The *in silico* analysis of the PbRpt3 protein identified different domains and motifs. In order to observe how they are displayed in the protein structure, a PbRpt3 3D model was predicted (Figure 1D) based on the crystallographic structure of the human Rpt3 (PSMC4 isoform 1 of 418 residues), as these two proteins share a sequence identity of 67% (Supplementary Figure 2). As expected, the 3D structure showed a coil region in its N-terminus, an OB-C-terminal domain (a.a. 64 to 119) including five stranded β-sheet coiled, with a very short α helix positioned between the third and fourth strands, followed by the ATPase domain comprising an α-β-α subdomain which includes the motifs involved in ATP binding and hydrolysis (the Walker A motif is located between a β-sheet and an α helix, and the Walker B motif forms a whole β-sheet). The AAA-lid domain is also present and reveals an all-α helix subdomain with 4 α helix between a.a. 320-360. Interestingly, our 3D model predicted that an Mg^2+^ ion would be involved in the stabilization of the PbRpt3 structure. The human Rpt3 (PSMC4) also possesses a Mg^2+^ ion [38]. By positioning the ATP on our 3D model, we observed that this molecule would mainly be maintained with hydrogen bonds located on the K188 and the Q356 of the protein (Figure 1E). N288 would also contribute to stabilize ATP through hydrogen bonds. In addition, ATP appears to be essential for the maintenance of the magnesium ion, and D241 would contribute to the stabilization of this ion. Thus, the 4 amino acids K188, D241, N288 and Q356 seem to play an important role in stabilizing the ATP molecule and, *in extenso*, they are important for the binding of Mg^2+^ and the structural arrangement of the protein. Of note, K188 and D241 are located in the Walker A and Walker B domains respectively (Figure 1C). Another part of the protein backbone of PbRpt3 would also contribute to the stabilization of ATP with the sequence 143-LGG-145. However, the mutation of these amino acids to alanine residues would not change the conformational structure of the pocket where the adenine group of ATP is located (data not shown).

### The two RVxF motifs of PbRpt3 are accessible for a possible interaction with PP1c

The 3D predicted model of PbRpt3 3D showed that the two RVxF motifs (200-KVTF-203 and 305-RKIEF-309) are located on either side of the protein, within the AAA-ATPase domain (Figure 1A, 1D). This accessible location would allow a possible interaction with PbPP1c. In the configuration shown, it seems unlikely that the PbPP1c protein could interact with both RVxF motifs at the same time. Indeed, the folding of PfPP1c has been modelled (https://swissmodel.expasy.org/repository/uniprot/Q8ILV1?csm=9CE0B7C29FEFEFC6), and it appears that its compact predicted conformation [39] wouldn’t allow the interaction with two RVXF motifs of PbRpt3 at the same time. However, we cannot exclude that the interaction between the two partners could involve a PbRpt3/PbPP1c protein ratio of 1:2.

### PbRpt3 would interact with PP1c independently of the proteasome complex

To better visualize the tightness of the packing around the modeled PbRpt3 protein in the proteasome complex, and to assess the accessibility of the RVxF motifs of PbRpt3 for a possible interaction with PbPP1c, we replaced the experimental human PSMC4 by the modeled 3D structure of PbRpt3 within the crystallographic structure of the human proteasome (RCSB ref. 6MSB). By analyzing this integration using the Chimera software, we observed, in the immediate environment of the RVxF motifs of PbRpt3, the B chain PSMC1 (Rpt2), the F chain PSMC3 (Rpt5), the K and L chains (alpha subunits) and the f chain PSMD2 (Rpn1). Within this chimera predicted complex, the B chain (Rpt2) appears to be able to contribute to the stabilization of the ATP of PbRpt3, and PbRpt3 would contribute to the stabilization of the ATP of the F chain (Rpt5). Thus, in this proteasome complex, we note that the KVTF and RKIEF motifs of PbRpt3 are not sufficiently accessible for a possible interaction with another protein than those mentioned. Should all the proteasome protein-protein interaction be conserved between Human and the *Plasmodium berghei* organism, the interaction between PbRpt3 and PbPP1 would therefore take place when PbRpt3 is untethered to the complex, implying a possible PbRpt3-PbPP1c interaction before the assembly and/or after the disassembly of the 26S proteasome complex.

### His-tagged PbRpt3 protein directly interacts with PP1c

The interactome of PbPP1c showed that PbRpt3 was identified in immunoprecipitated protein complexes by spectrometry analysis [15,16]. In order to investigate whether these proteins interact directly, we produced a recombinant protein corresponding to the PbRpt3 full length (aa 1-395, 45kDa, Supplementary Figure 3A) and performed an ELISA-based assay using biotinylated PP1c and PfI2 as a positive control [33]. The results shown in Figure 2A demonstrate that PbRpt3-45 kDa directly interacts with PfPP1c, as evidenced by the dose-dependent increase in OD when PbRpt3 is incubated with biotinylated PP1c. To determine further which region of PbRpt3 interacts with PP1c, we produced a recombinant truncated PbRpt3 (aa 131-395, 30 kDa, Supplementary Figure 3B). This fragment lacks the coil region and the ATPase OB-C-terminal domain. Interestingly, this truncated PbRpt3 also interacts with PfPP1c in a concentration-dependent manner in ELISA-based assay (Figure 2A). Taken together, these results suggest that the PP1c-interaction domains of PbRpt3 may lie in this region where both RVxF motifs are present.

**FIGURE 2.**
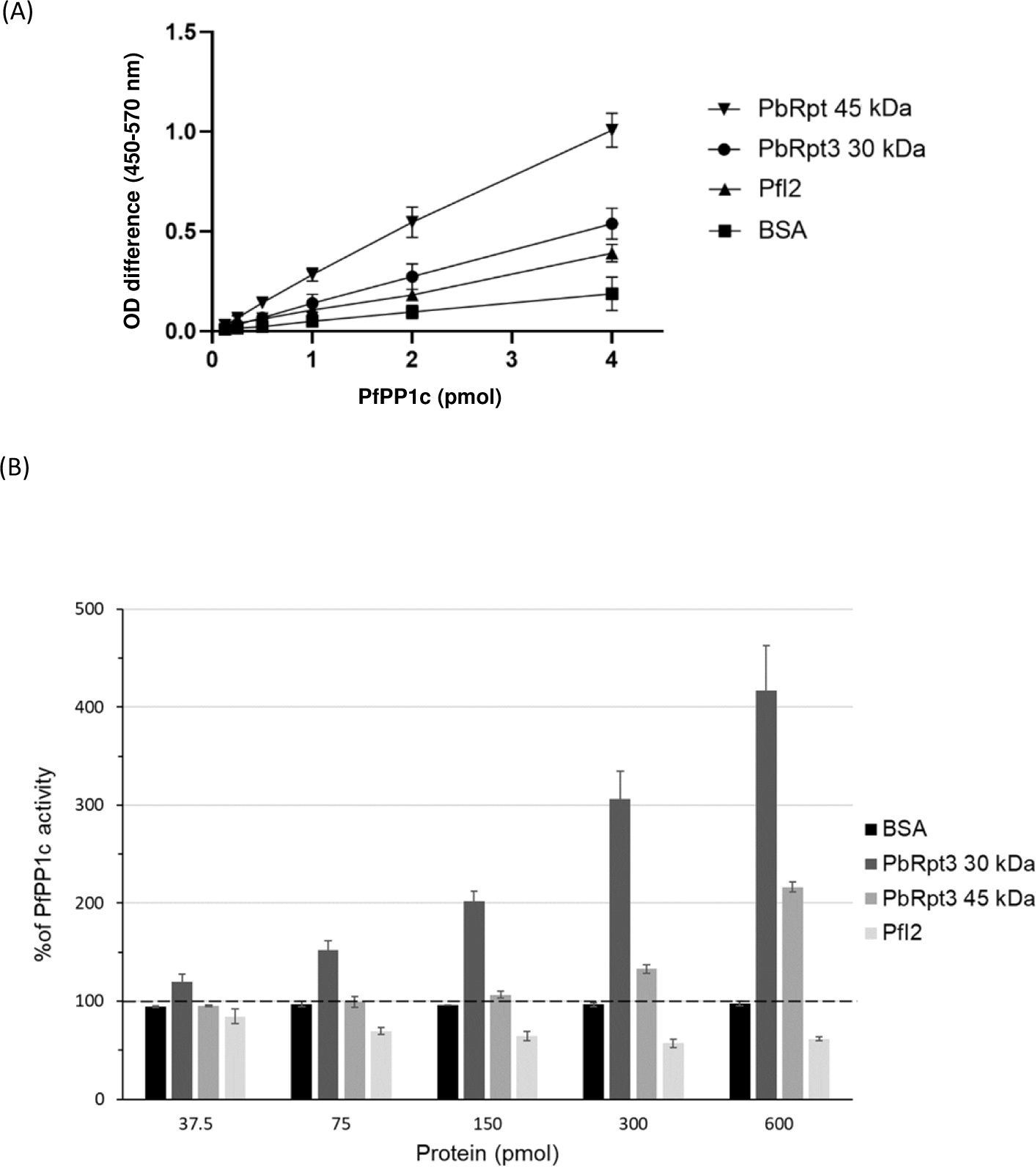
Interaction of PbRpt3 with PP1c in vitro and effect on phosphatase activity. **A**. Interaction study of PbRpt3-45 kDa and PbRpt3-30 kDa with PfPP1c using PfI2 as a positive control and BSA as a negative control. Results are representative of two independent experiments. **B.** Effect of PbRpt3-45 kDa, PbRpt3-30 kDa, BSA and PfI2 on PfPP1c phosphatase activity. Different quantities of each protein (37.5 pmol, 75 pmol, 150 pmol, 300 pmol and 600 pmol) were incubated with 1µg of recombinant PfPP1c for 30 min before the incubation with the substrate pNPP for 1 hour at 37°C. Results presented as % of relative increase or decrease of phosphatase activity are means ± S.D. of two independent experiments performed in duplicate.

### Effect of PbRpt3 on phosphatase activity of PP1c

As PbRpt3 recombinant protein interacts with PP1c, we assessed its effect on the phosphatase activity using the p-Nitrophenyl phosphate (pNPP) as a substrate. First, a control experience showed that when pNPP was used as substrate in the presence of PbRpt3 alone (without PfPP1c), no phosphatase activity could be detected (data not shown). In the presence of PfPP1c, the recombinant protein PbRpt3-45 kDa strongly increased the dephosphorylation activity of the phosphatase in a concentration dependent manner (Figure 2B) with a mean value of 220 % of PP1c activity when a quantity of 600 pmol of PbRpt3 was used. Similarly, the PbRpt3-30 kDa recombinant protein also increased PfPP1c activity, reaching more than 400% PP1c activity when incubating with 600 pmol of the truncated PbRpt3 (Figure 2B). As expected, PfI2, a known inhibitor of PP1c activity [40], decreased PfPP1c activity (∼60% of PfPP1c activity using 600 pmol of PfI2). These results strongly suggest that PbRpt3 would be an activator of PP1c. Moreover, although we cannot compare the values obtained with PbRpt3-45kDa and PbRpt3-30kDa due to a potential difference in the purity of the two recombinant proteins, these results clearly indicate that the activity of PbRpt3 on PfPP1c is carried by the aa 131-395 region of the protein.

### PbRpt3 binds to PP1c and shows functional activity in *Xenopus* oocytes

To further determine the functional activity of PbRpt3 in a cellular context, we took advantage of the *Xenopus* oocyte model in which it has been previously shown that several phosphatase partners could regulate cell-cycle progression from G2 to M, assessed by the appearance of GVBD (Germinal Vesicle Break Down) [32,35,40,41]. This is based on the fact that *Xenopus laevis* PP1c (XePP1c) shows >80% identity at the a.a. sequence level with PbPP1c [35]. In the *Xenopus* model, immature oocytes are blocked in prophase I and the inhibition of PP1c by anti-PP1c antibodies or by an inhibitor of this phosphatase will trigger the GVBD reflecting the G2/M transition to metaphase II [35,40–42]. Conversely, a PP1c activator will lead to an inhibition of the progesterone (PG)-induced maturation. To confirm the *in vitro* results showing the activation of PP1c by the recombinant PbRpt3, we microinjected the cRNA coding for the HA-tagged PbRpt3 protein (Supplementary Figure 4). First, the efficient translation of PbRpt3 in oocytes after cRNA microinjection was checked by immunoblot using anti-HA mAb (Supplementary Figure 5). Next, we performed immunoprecipitations of oocyte extracts using anti-*Xenopus* PP1c (XePP1c) mAb. The immunoblot analysis revealed that PbRpt3 and XePP1c were present in the same complex (Figure 3A). At the functional level, the microinjection of PbRpt3 cRNA alone did not induce the maturation of *Xenopus* oocyte, while PG or PfI2 did induce GVBD (Figure 3B). When oocytes maturation was induced with PG, we observed that the microinjection of PbRpt3 cRNA resulted in a significant reduction of the percentage of GVBD (Figure 3B; mean percentage of GVBD : 86.7% and 1.7% for PG and PG+PbRpt3 cRNA respectively). Taken together, these results show that PbRpt3 seems to activate XePP1c in oocytes, confirming the results obtained *in vitro*.

**FIGURE 3.**
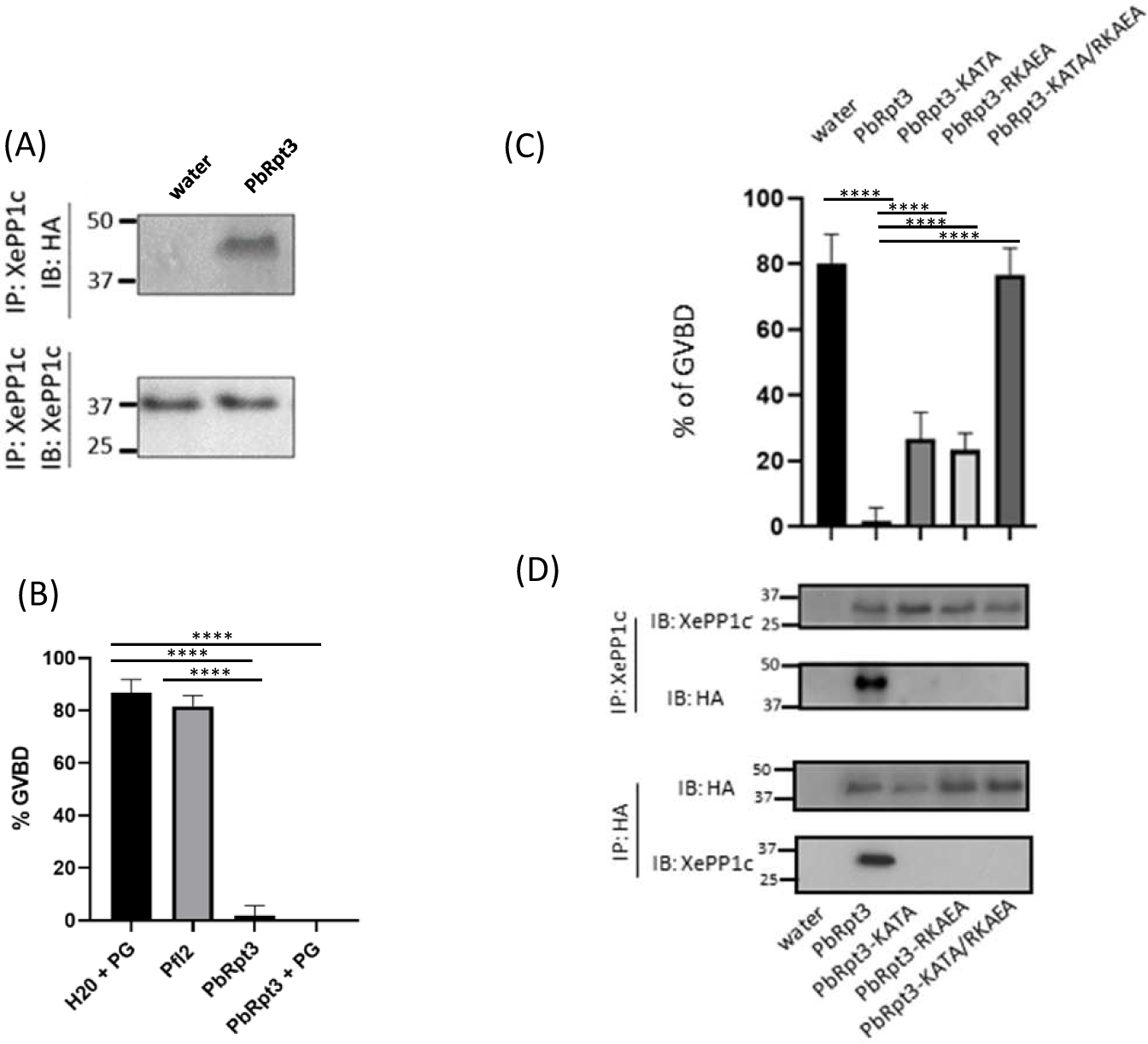
PbRpt3 functional analysis using *Xenopus* oocytes model. **A.** Interaction of PbRpt3 with *Xenopus* PP1c (XePP1c). Extracts prepared from *Xenopus* oocytes previously micro-injected with either water (80 nl, lane 1) or PbRpt3 cRNA (60 ng in 80nl, lane 2), followed by progesterone (PG) treatment (10 μM, 15 hours after micro-injection) were immunoprecipitated using an anti-XePP1c mAb. An immunoblot analysis was performed using anti-HA mAb (upper panel) and anti-XePP1c mAb (lower panel). **B, C.** Role of PbRpt3 on the induction of Germinal Vesicle BreakDown (GVBD). Values are presented as mean percentages, and p value <0,0001 for each test. Each experiment was performed using a set of 10 oocytes and repeated on three animals. **B.** The appearance of GVBD induced by PG 15 h after microinjection of water (60 nl as control, lane 1) is strongly inhibited by the micro-injection of PbRpt3 cRNA (20 ng in 60 nl, lane 4). In the absence of PG, the micro-injection of PbRpt3 cRNA (20 ng in 60 nl, lane 3) is not sufficient to trigger the GVBD. As a control, the microinjection of the PP1c inhibitor PfI2 cRNA (20 ng in 60 nl, lane 2) triggers the oocytes GVBD. **C.** Role of the RVxF motifs of PbRpt3 for its functional activity. Percentage of GVBD of oocytes which have been micro-injected with either water as control (80 nl, lane 1), or cRNAs (60 ng in 80 nl) coding for PbRpt3 (lane 2), PbRpt3-KATA (lane 3), PbRpt3-RKAEA (lane 4) or PbRpt3-KATA/RKAEA (lane 5). All micro-injections were followed by PG treatment (10 μM, 15 hours after micro-injection). **D.** Role of the RVxF motifs of PbRpt3 for its interaction with XePP1c. Oocytes were micro-injected with water as control (80 nl, lane 1), or cRNAs (60 ng in 80 nl) coding for PbRpt3 (lane 2), PbRpt3-KATA (lane 3), PbRpt3-RKAEA (lane 4), or PbRpt3-KATA/RKAEA (lane 5), 15 hours prior incubation with PG (10 μM). Then, the extracts were immunoprecipitated with anti-XePP1c mAb (upper panel) or anti-HA mAb (lower panel) and subjected to immunodetection using the same antibodies. Each experiment was performed using a set of 20 oocytes and repeated on two animals.

### PbRpt3 RVxF motifs are crucial for PP1c binding and functional activity in *Xenopus* oocytes

Many studies showed that the binding of PP1c to RVxF-dependent interacting proteins could be disrupted when their RVxF amino acids V/I or F are substituted with alanine residues [43,44]. In order to explore the contribution of RVxF motifs of PbRpt3 in its functional activity, we generated PbRpt3 cRNAs bearing single-motif mutations either on 200-KVTF-203 (PbRpt3-KATA) or 305-RKIEF-309 (PbRpt3-RKAEA), or double-motif mutations (PbRpt3-KATA/RKAEA) (Supplementary Figure 4). The effective translation in the oocytes of each microinjected cRNA was checked by immunoblot using anti HA mAb (Supplementary Figure 5). We then assessed the capacity of each single mutated protein as well as the double-mutated protein to inhibit the PG-induced GVBD. As with PbRpt3, the microinjection of either single mutated cRNA, PbRpt3-KATA or PbRpt3-RKAEA, resulted in a significant reduction in the PG-induced GVBD (Figure 3C, lanes 3 and 4). However, this inhibition seemed to be partial when single mutated cRNAs were used (mean percentage of GVBD of 80%, 1.7%, 26.6%, and 23.3% for control, PbRpt3, PbRpt3-KATA or PbRpt3-RKAEA respectively). The co immunoprecipitation/immunoblot assays performed on lysates prepared from these oocytes indicated that the PbRpt3-XePP1c complex was not detected when single mutants were injected (Figure 3D). The partial functional effect of single mutants may thus be attributed to a residual and/or transitory binding of PbRpt3 to XePP1c that would not be detected by immunoblot. In addition, and as mentioned above, a possible PbRpt3/PP1c ratio of 1:2 could also explain the limit of detection of PP1, due to the lower level of XePP1c in the complex. To further define the contribution of both RVxF motifs, we microinjected the double mutated cRNA (PbRpt3-KATA/RKAEA) in the oocytes and followed up the appearance of GVBD. As shown in Figure 3C, this double mutant was not able to inhibit PG induced GVBD (lane 5; percentage of GVBD 76%). In addition, PbRpt3-KATA/RKAEA mutant protein failed to co-immunoprecipitate with XePP1c (Figure 3D, lane 5), despite its effective translation by the oocyte (Supplementary Figure 5, lane 6). Taken together, these results demonstrate that PbRpt3 directly interacts with PP1c, and that its interaction with PP1c *via* both RVxF motifs would be required for its functional effect.

### The ATPase function of PbRpt3 is involved in the activation of XePP1c

PbRpt3 is expected to be an AAA-ATPase, as evidenced by the domain present in its protein sequence. In the predicted 3D structure, we observed that 4 amino acids (K188, D241, N288, and Q356) are involved in stabilizing the ATP molecule, as well as in the binding of a Mg^2+^ion. The mutation of these amino acids may thus disrupt the enzymatic activity of the protein. To explore whether the ATP binding of PbRpt3 is involved in its functional activity towards PP1c in *Xenopus* oocytes, we generated a mutated cRNA for PbRpt3 where all the 4 residues K188, D241, N288 and Q356 (Supplementary Figure 4) were replaced by an alanine. The results in Figure 4 showed that the mutation of these 4 amino acids partially abolished the inhibition of PG-induced GVBD by PbRpt3. The microinjection of the PbRpt3 cRNA mutated for the ATP-binding sites resulted in a 39.2 % GVBD, while PbRpt3 cRNA microinjection resulted in 1.7% GVBD (mean percentage of GVBD in PG-treated oocytes: 80.8%). This experiment was repeated using 3 different batches of PbRpt3 cRNA mutated on the 4 ATP-binding sites and a total of 100 *Xenopus* oocytes and the same partial effect was observed in each case, with a GVBD only reaching a mean percentage of 40% indicating that these results are reproducible and accurate. Interestingly, we observed that in the subsequent immunoprecipitation experiments, the mutated PbRpt3 protein conserved its binding capacity to XePP1c (Figure 4B, lane 3). The partial effect observed on PG-induced GVBD could be explained either by a partial contribution of the ATPase activity of PbRPt3, or by a possible remaining ATPase activity related to a residual capacity of the mutated protein to bind an ATP molecule. In fact, as described above, the sequence 143-LGG-145 may be involved in stabilizing the ATP even in the absence of certain crucial residues granting the presence of stabilizing hydrogen bounds. However, as described earlier, structure modelling predicted that mutating these three amino acids by alanine residues would bring a direct effect neither on the interaction of the lateral chains with ATP, nor on the backbone organization of this part of the protein, thus predicting a high stability of the ATP maintaining pocket. Taken together, these findings indicate for the first time that the ATPase activity of PbRpt3 could contribute to the activation role of PbRpt3 towards the phosphatase PP1c.

**FIGURE 4.**
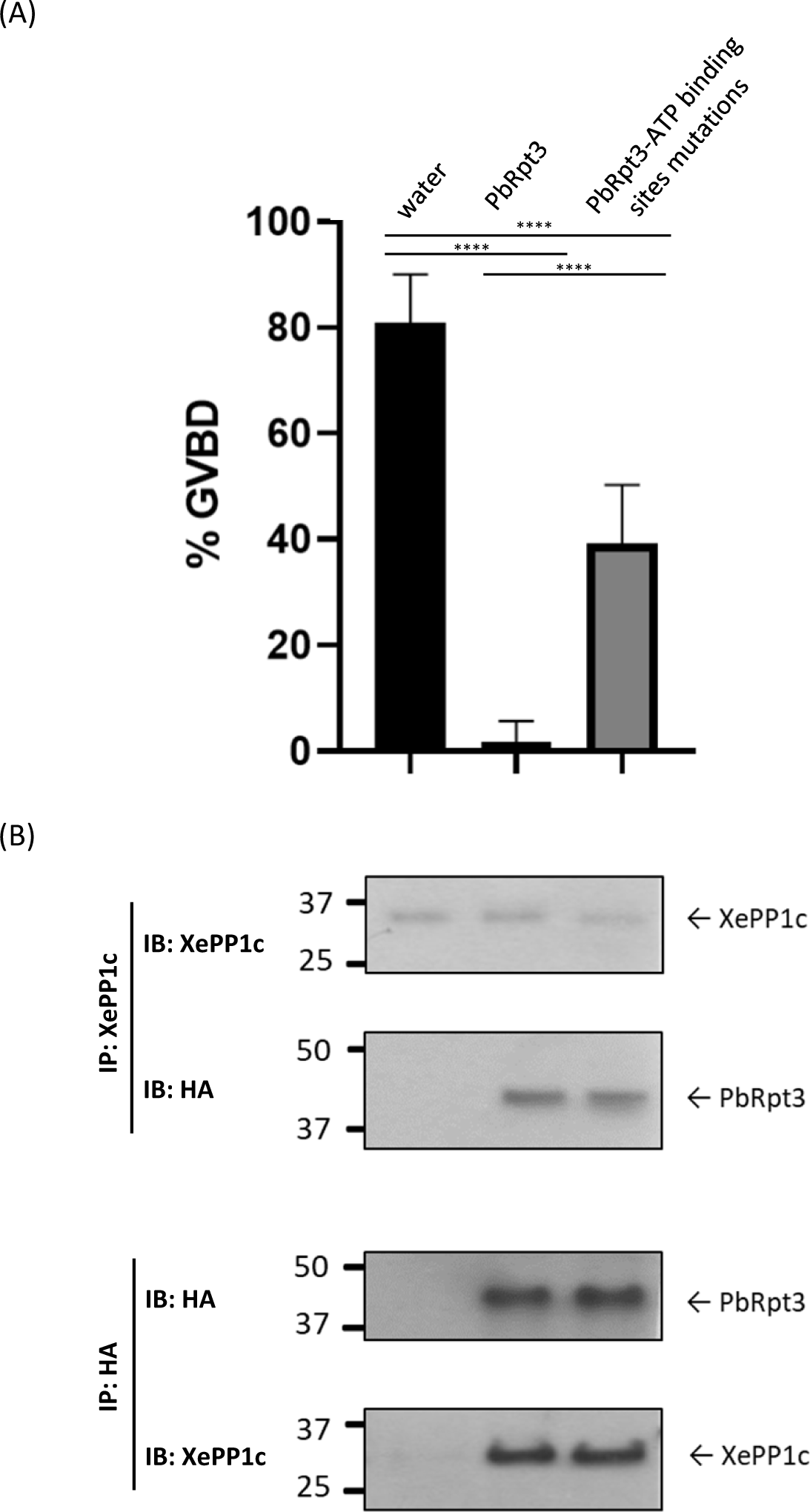
Role of the predicted ATP-binding sites of PbRpt3 in its functional activity in *Xenopus* oocytes. **A.** Percentage of GVBD of oocytes which have been micro-injected with either water as control (80 nl, lane 1), PbRpt3 cRNA (60 ng in 80 nl, lane 2) and PbRpt3 cRNA mutated on 4 ATP binding sites (60 ng in 80 nl, lane 3). All micro-injections were followed by progesterone treatment (10 μM, 15 hours after micro-injection) and the graph is representative of 40 oocytes from two independent animals. Values are presented as mean percentages, and p value <0,0001 for each test. **B.** Interaction between XePP1c and PbRpt3 analyzed by immunoprecipitation. 20 oocytes were micro-injected with water as control (80 nl, lane 1), PbRpt3 cRNA (60 ng in 80 nl, lane 2) and PbRpt3 cRNA mutated on 4 ATP binding sites (60 ng in 80 nl, lane 3), 15 hours prior incubation with PG (10 μM). The extracts were immunoprecipitated with anti-XePP1c mAb (upper panel) or anti-HA mAb (lower panel) and subjected to immunodetection using the same mAbs. Each experiment was repeated on two animals.

### PbRpt3 may be essential during the blood stage life cycle of *P. berghei*

Our results obtained *in vitro* and in *Xenopus* oocytes demonstrated an interaction of PbRpt3 with PP1 catalytic subunit, as well as a regulatory role of the AAA-ATPase -partially *via* its ATPase function on the dephosphorylation activity of PP1c. In an attempt to understand the role of Rpt3 in the parasite, we used reverse genetics in *P. berghei*. In a previous *P. berghei* high throughput functional analysis of disruption phenotypes, PbRpt3-disrupted mutants showed a ‘slow’ growing rate phenotype in mice, while 4 out of the 5 other Rpt proteins were suggested to be essential for the erythrocytic cycle [45]. To further study whether the lack of PbRpt3 could affect *P. berghei* blood stages life cycle, we attempted to generate knock-out (KO) lines by taking advantage of the available PlasmoGEM plasmid (PbGEM-022521; 93 % deletion of the PbRpt3 gene). Transfections were performed in two different strains of *P. berghei* ANKA (pG230 and PbGFP) with this KOgem construct. After selection with pyrimethamine, resistant parasites were genotyped by PCR (See Supplementary Table 2 and Supplementary Figure 6A). A total of 4 independent transfections were performed, allowing the detection of parasites 7 to 9 days after transfection under pyrimethamine selection. In two of these transfections, the diagnosis genotyping showed either no integration of the resistance cassette at the PbRpt3 locus, or an integration only of the 5’ side. After two other transfections, we detected both the integration of the construct on 5’ and 3’ sides, and the presence of the wild-type PbRpt3 gene (Supplementary Figure 6B). In order to increase the ratio of transgenic *vs* wild-type parasites, we first performed up to 7 successive passages of the parasites in mice under pyrimethamine regimen. However, this did not lead to any enrichment in transgenic parasites (data not shown). Next, a total of five attempts were performed to clone the parasites by limiting dilution either from the newly resistant parasites (first appearance after transfection and drug selection) or from different passages under pyrimethamine pression. Depending on the quantity of parasites injected per mouse, we either obtained no parasites in the recipient mice (3 experiments with a total of 26 recipient mice), or parasites showing both wild-type and KO construct-integrated genotype (for 1 experiment, 2/10 mice have shown parasites). In the fifth cloning experiment (20 parasites/mouse, 10 recipient mice), 2/10 mice showed blood parasites, and after passage on 3 mice, the parasites were genotyped as pure wild type parasites (Supplementary Figure 6C). In summary, all attempts to select or clone parasites which have integrated the dhfr resistance cassette at the *PbRpt3* locus were either unsuccessful or only allowed to clone wild-type parasites. These results are supportive of the essentiality of PbRpt3 during the erythrocytic stage of *P. berghei*. This could be interpreted as differing from the results previously obtained by Bushell et al showing a slow growing rate of PbRpt3-KO parasites at erythrocytic stages [45]. However, in this global analysis, mutants were generated by simultaneous co-transfection of multiple barcode vectors, which would help explain this difference. In *P. falciparum*, a saturation mutagenesis study showed that most of the proteasome 26S genes, including PfRpt3, were essential for parasite survival [46]. In addition, selective inhibitors of the catalytic subunits of the parasite proteasome have shown potent anti-malarial effect lethal effect [17,18,47]. Therefore, PbRpt3 essentiality may be related either because of its role into the proteasome complex, or independently of the proteasome, complexed to different partners and throughout other pathways.

### Localization of PbRpt3 in *Plasmodium berghei*

In order to localize PbRpt3 throughout the erythrocytic cycle of the parasite, we generated parasites expressing mCherry-tagged PbRpt3 using a single homologous strategy (Figure 5A). The correct integration of the mCherry tag was checked by PCR genotyping (Figure 5B). Following enrichment of the mCherry positive parasites by cell sorting, the expression of the tagged protein was checked by immunoblot analysis (Figure 5C). PbRpt3 localization was then assessed by immunofluorescence assays on different erythrocytic stages of *P. berghei*. Figure 5D shows the expression of PbRpt3 during the asexual development, with a low expression in early trophozoite stages which strongly increases during the late trophozoite stages. In schizonts, the signal corresponding to PbRpt3-mCherry seems to be less intense. Gametocytes show also a high level of PbRpt3-mCherry expression. This expression profile is consistent with the results of transcriptomic studies showing a maximum transcript level at the trophozoite and schizont stages [48]. In addition, the absence of the transcript at the gametocyte stage, combined with our observations, could indicate storage of the protein at this stage. Concerning the cellular distribution of PbRpt3-mCherry, we observed that it is mainly cytoplasmic in trophozoites and schizonts stages, with a punctuated localization particularly observed in late trophozoites. This localization may correspond to parasites organelle structures or to proteasome storage granules. The latter have been described in the cytoplasm of quiescent yeasts [49]. In gametocytes, the IFA signal overlaps with DAPI staining, indicating that PbRpt3 is more accumulated in the nucleus at this stage. Overall, the distribution of PbRpt3 fits with the localization of the proteasomes in eukaryotic cells as they are mainly found in the cytoplasm of the cell, associated with the centrosomes, the cytoskeletal networks and the outer surface of the endoplasmic reticulum [3,4]. However, proteasomes have also been described in the nuclei of eukaryotic cells where they are present throughout the nucleoplasm. But their relative abundance within nuclei and cytoplasm compartments is variable, often depending on the cell type, growth status and stimulation or stress conditions.

**FIGURE 5.**
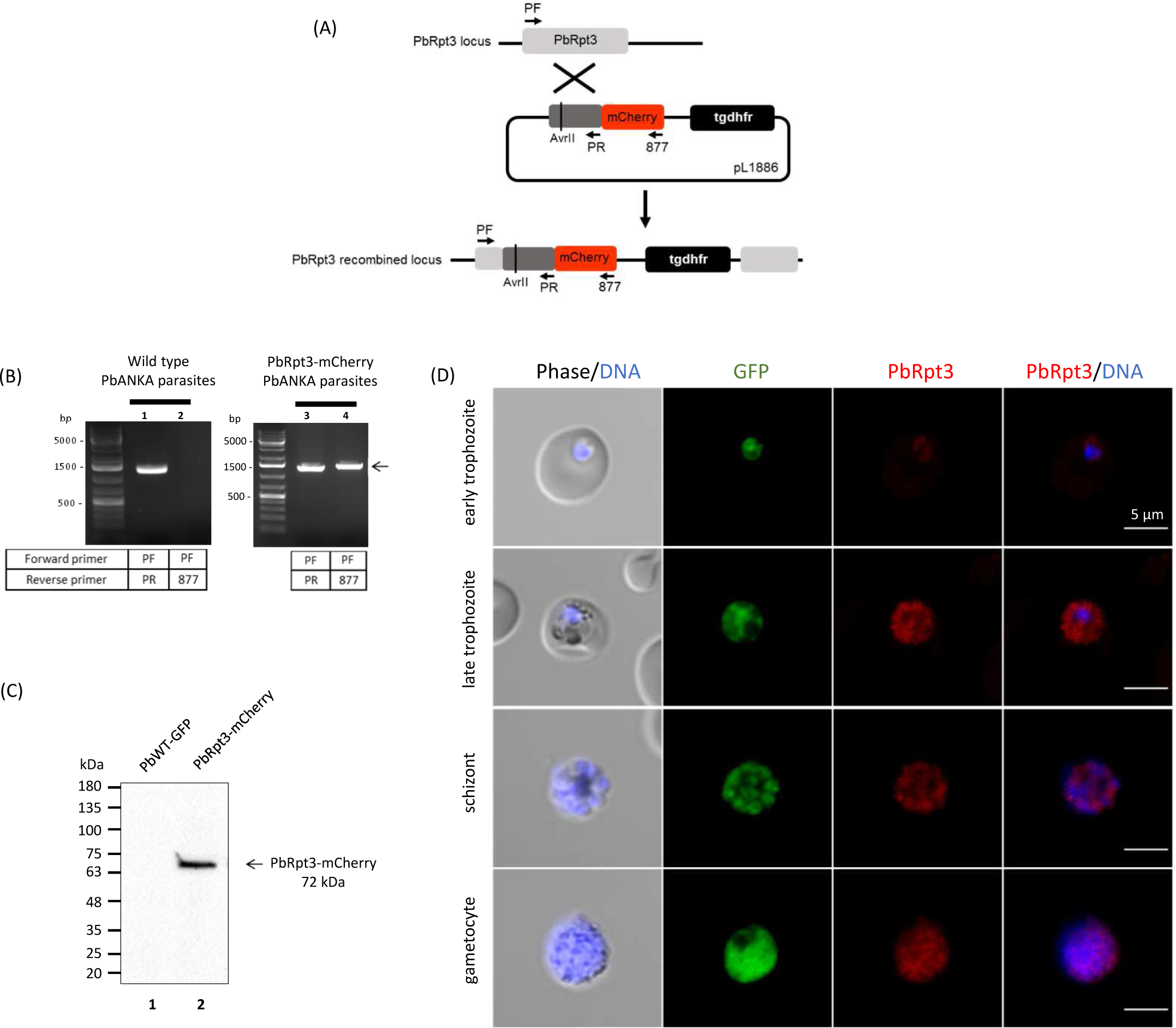
Tagging of endogenous Rpt3 in *Plasmodium berghei* and localization studies A. Knock-in strategy using the vector pL1886 that allows the insertion of the mCherry epitope at the C-terminus of PbRpt3 with a single homologous recombination. The pL1886 construct, the tgdhfr-resistance cassette, the location of the primers used for PCR analysis and the locus resulting from integration are shown. A silent mutation was inserted in the coding sequence to generate an AvrII single cutter restriction site, in order to linearize the plasmid before transfection into PbANKA-GFP strain. **B.** Diagnosis PCR analysis of pL1886-tgdhfr-PbRpt3-mCherry transfected *Plasmodium berghei* parasites. Lanes 1 and 2 correspond to DNA extracted from wild-type (WT) *P. berghei* ANKA parasite and lanes 3 and 4 correspond to DNA extracted from transfected parasites. Lanes 1 and 3 represent the detection of the wild-type locus (PCR with PF and PR); and lanes 2 and 4 correspond to the detection of the 5’ integration at the PbRpt3 locus (PCR with PF-877, arrow). **C.** Immunoblot analysis of pL1886-PbRpt3-mCherry transfected *P. berghei*. Proteins extracted from wild type parasites (lane 1) or from transfected parasites (lane 2) were subjected to western blotting, probed with anti-RFP antibodies. **D.** Cellular distribution of PbRpt3-mCherry in *P. berghei* erythrocytic stages (early trophozoite, late trophozoite, schizont and gametocyte), analyzed by immunofluorescence with anti-mCherry antibodies. The PbRpt3 protein appears in red and the DNA is stained with DAPI. The parasites express the Green Fluorescent Protein (GFP).

### Analysis of PbRpt3 interactome: the evidenced presence of the 19S proteasome complex

In order to explore the pathways involving PbRpt3 in *Plasmodium*, we performed a global immunoprecipitation of PbRpt3-mCherry obtained from soluble extracts of a mixed schizonts population (early, middle and late), followed by a mass-spectrometry analysis of the protein complexes (IP/MS). Four biological replicates were analysed by IP/MS along with wild-type parental line used as a negative control for contaminant interactions. This analysis led to the identification of 2713 proteins, of which 623 were significantly enriched in the PbRpt3-mCherry IP (T test q-value < 0.01). (Supplementary Table 3, sheet 1). The detection of PbRpt3 bait confirmed the quality of the IP/MS. The entire putative proteasome regulatory particle ATPase Rpt (Rpt1 to 6) were identified, as well as all the Regulatory particles non-ATPase Rpn (Rpn 1 to 13) (Figure 6A). Of note, none of the alpha or beta subunits belonging to the catalytic 20S proteasome complex were present among the significantly enriched 623 proteins. In addition, no recovery of mouse 20S subunits was detected (data not shown). A previous study reported the characterization of the 26S proteasome of *P. falciparum*, using an affinity purification strategy based on the ubiquitin-like domain of PfRad23, one of the ubiquitin receptors which target ubiquitylated proteins to the proteasome [20]. In this study, all the 19S and 20S proteasome subunits were identified in a mixed population of trophozoites and early schizonts. However, *P. falciparum* parasites were subjected to formaldehyde crosslinking prior to analysis, thus allowing to detect both stable and transient interactions. Especially, this procedure allowed to recover 20S subunits of the proteasomes. As mentioned by the authors, these 20S subunits were only present in very low quantities in the non-crosslinked control samples. These observations, together with the absence of the *P. berghei* 20S subunits in our study, suggest that the association of 19S and 20S is transient and dynamic in the parasite.

**FIGURE 6.**
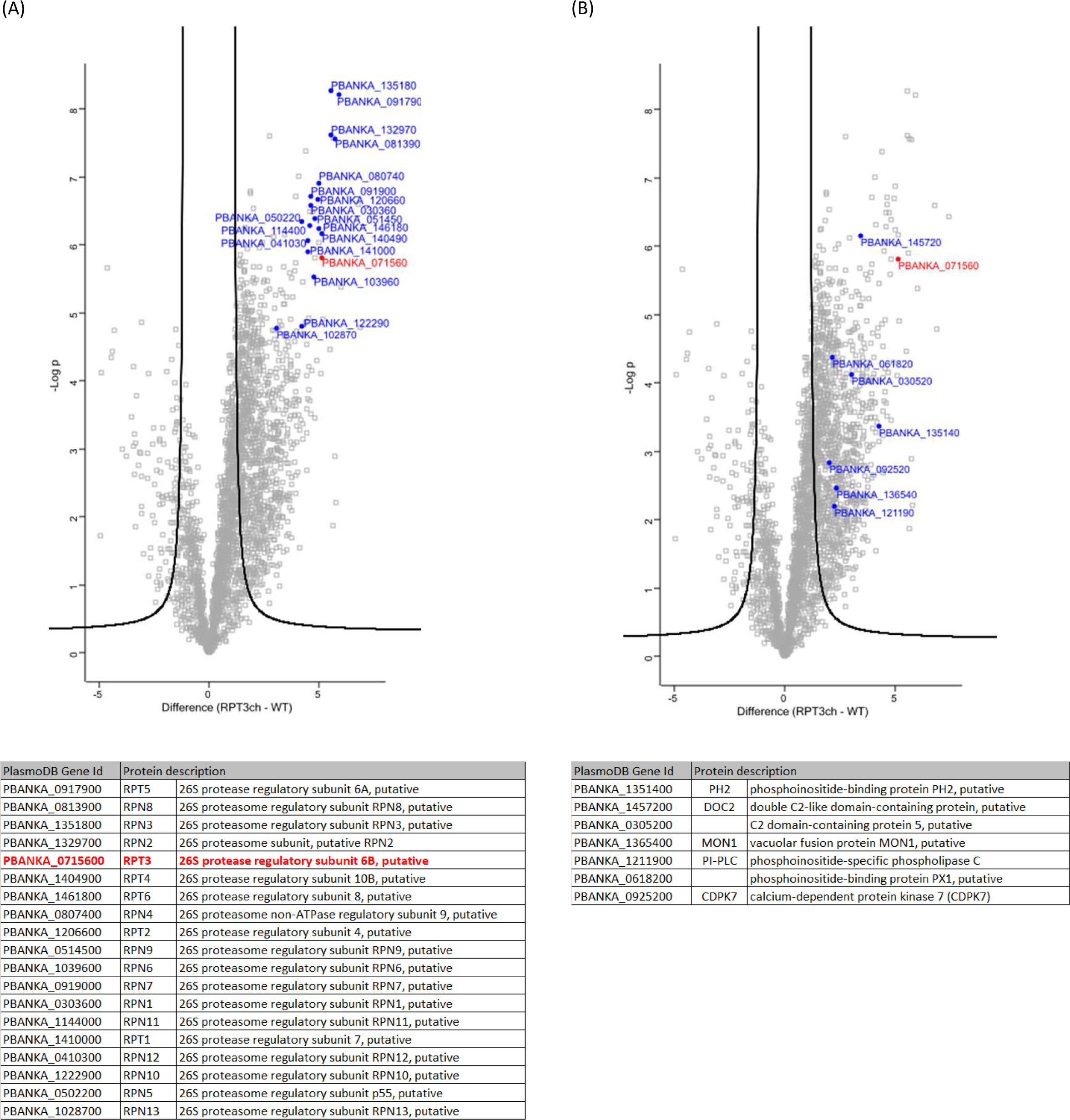
PbRpt3 interactome analysis. Volcano plot representations of the outcome of the PbRpt3 interactome study. PbRpt3 (PBANKA_0715600) is highlighted in red. Highlighted in blue are the interacting proteins belonging to the 19S proteasome (**A**) and to the GO enriched “phospholipid binding pathway” (**B**). List of the proteins are included in the tables below the plots, ranked according to their Student’s t-test difference. Gene identifiers (Id) and protein descriptions were updated using PlasmoDB relapse 64 (12 jul 2023), except for RPN4 and RPT2 which short names have been attributed based on sequence homology.

As mentioned above, the 3D model suggested that PP1c would not be able to interact with PbRpt3 when complexed in the 26S proteasome. Thus, with the absence of the 20S proteasome subunits from our interactome study, it could be expected to detect PP1c in the protein complexes. However, PbPP1c was not detected among the 623 PbRpt3 potential partners. PbRpt3 was initially identified in PbPP1c interactome, both in schizonts and gametocyte stages [16]. However, we have already observed that PP1c is not always detected in IP/MS analyses using PP1c partners as bait, despite direct interaction *in vitro* and detection of the complex in the parasite [15]. Interestingly, 6 other phosphatases, and 20 kinases are identified among the 623 PbRpt3 potential partners (Supplementary Table 3, sheet 1). These include Casein kinase 2 (CK2), which has already been shown to phosphorylate 20S proteasome subunits in eukaryotes [50]. These observations suggest that phosphorylation processes may be involved in the assembly and/or regulation of the proteasome 19S in *Plasmodium*.

To further gain new insights into pathways involving PbRpt3 in the parasite, we performed a Gene Ontology enrichment analysis including the 623 potential partners using PlasmoDB software (Release 63, May 2023). Enriched GO terms/pathways of computed and curated biological processes, cellular components and molecular functions were analysed and summarized in Supplementary Table 3, sheet 2. Ten GO Terms were found significantly enriched, corresponding to 39 different *P. berghei* proteins, including PbRpt3 (Supplementary Table 3, sheet 3). Among them, 19 proteins belong to the Ubiquitin proteasome pathway. This includes the 6 putative proteasome regulatory particles ATPase Rpt (Rpt1 to 6), as well as the 13 Regulatory particles non-ATPase Rpn (Rpn 1 to 13). We did not observe the presence of other proteasome-related proteins such as the two proteasome ubiquitin receptors, Rad23 and Dsk2 which have been identified in the analysis performed by Wang et al. [20].

### PbRpt3 interaction network includes proteins related to gene transcription and phospholipids binding proteins

Beside the 26S proteasome-related pathways, our GO analysis revealed a significant enrichment in “intracellular protein-containing complex” pathway (Cellular components, GO:0140535) and “phospholipid binding pathway” (Molecular function, GO:0005543) (Supplementary Table 3, sheet 2). In order to get insights into the potential relation with the proteasome 19S subunit, the 39 proteins belonging to enriched pathways underwent the retrieval of networks for interacting proteins using STRING software (v.11.5, July 2023). Interestingly, proteins known to regulate gene transcription were found in this protein–protein interaction network (Supplementary Figure 7). This includes the two acetyltransferases MYST (PBANKA_0929500) and ARD1 (PBANKA_1201700) as well as the DNA directed RNA polymerase 3 largest subunit (PBANKA_1344000) and the CDC73 domain-containing protein (PBANKA_1235400). This observation, together with the presence of the transcription factor TFIID (PBANKA_0829800) in the list of the 39 proteins (not present in the predicted interaction network), suggest that PbRpt3 and/or the 19S regulatory particle of the proteasome may be linked to transcription regulation in the parasite. Our GO enrichment and STRING analyses of PbRpt3 interactome has also uncovered 7 proteins belonging to the “phospholipid binding pathway” (Figure 6B, Supplementary Table 3, sheet 3; Supplementary Figure 7). Among these proteins, we find the phosphoinositide-binding protein PH2 (PBANKA_1351400), which has been described to play a role during the fusion between the micronemes membrane and cytoplasmic membrane in *P. falciparum* [51], the phosphoinositide-specific phospholipase C (PI-PLC, PBANKA_1211900), the double C2-like domain-containing protein (DOC2, PBANKA_1457200) and the phosphoinositide-binding protein (PBANKA_0618200). We also observe, linked to this last protein in our STRING analysis, the calcium dependent protein kinase 7 (CDPK7, PBANKA_0925200). Interestingly, PfCDPK7 interacts with 4’ phosphorylated phosphoinositides and is involved in the regulation of phospholipid biosynthesis in *P. falciparum* [52]. Moreover, PfCDPK7 was described to be a partner of PfRCC-PIP, a *Plasmodium*-specific regulator of PfPP1c, suggesting that PfRCC-PIP could provide a platform to regulate phosphorylation / dephosphorylation processes [35]. Taken together, although these potential interactions should be confirmed by complementary approaches, our observations support a role for PbRpt3, within the 19S proteasome or alone, in various crucial parasite mechanisms via its binding to phospholipids.

## Materials and Methods

### Ethics statement

Mice were purchased from Charles River and housed in an Animal Biosafety Level 2 facility at the Institut Pasteur de Lille. Mature *Xenopus laevis* females were purchased from the CRB-University of Rennes I, and housed in PHExMAR, University of Lille. All animals were maintained in accordance with the French National Guidelines for Use of Animals for Scientific Purposes which is also in line with EU Directive 2010/63/EU. Experimental protocols performed in this study were reviewed and approved by the “Comité d’Ethique CEEA-75 en Experimentation Animale Nord-Pas de Calais-France” (mice project number: 18905-2019020111166978v2; *Xenopus* project number: F59-00913).

### Animals

Mice: Infections with the parasite *Plasmodium berghei* were performed in 4 weeks CD1 male mice maintained in filter cages and sorted randomly into groups of 3-4 animals. Mice were infected with parasites resuspended in phosphate buffered saline by intraperitoneal injection. Drugs were administered in drinking water or by intraperitoneal injection. Intra-erythrocytic parasitemia was monitored regularly on blood smears.

*Xenopus*: After anesthesia, realized by immersion in 2 g/L MS222 solution (tricaine methane sulfonate), ovarian lobes were surgically removed and stored in ND96 medium (96 mM NaCl, 2 mM KCl, 1.8 mM CaCl2, 1 mM MgCl2, 5 mM HEPES-NaOH, pH 7.5) at 19 °C.

### PbRpt3 sequence analysis and molecular modeling

We used PlasmoDB data base version 63 as a reference for gene annotation and expression [https://plasmodb.org]. The model was constructed using the sequence alignment of *Plasmodium berghei* PbRpt3 <u>[PBANKA_0715600]</u> and the human PSMC4 of 418 amino acids (PDB file 6MSB – chain D). This protein shows 67% identity with PbRpt3 (Supplementary Figure 2). The crystallographic structures available on PDB (Protein Data Bank, http://www.rcsb.org/pdb/) allowed us to build a 3D model of PbRpt3 by homology modeling. The magnesium ion was copied from the reference, the atomic partial charges calculated with the Gasteiger-Hückel method, the geometry of the backbone was optimized by 2000 iteration steps of conjugated gradients, and the protein was allowed to move until the gradient value was smaller than 0.01 kcal mol^−1^ Å^−1^ with the MMFF94s force field [53]. The quality of the model was checked with its Ramachandran plot (Supplementary Figure 8). The calculations were performed with SYBYL software version 6.9.2 (Tripos Associates, St. Louis, Missouri) and the images with Chimera software version 1.15. PbRpt3 was finally placed in the human proteasome by replacing the D chain in order to have a representation of its environment (PDB file 6MSB).

### Plasmids and directed mutagenesis

Plasmids pGADT7 and pETDuet-1 were purchased from Clontech and Novagen respectively. The plasmid pL1886 was kindly provided by Dr B. Franke-Fayard (Leiden University Medical Center, The Netherlands). The plasmid PbGEM-022521 used in the reverse genetic study in *P. berghei* was kindly gifted from the Plasmogem-Wellcome Sanger Institute (Cambridge, UK). For *Xenopus* oocytes experiments, a PbRpt3 cDNA with optimized codons has been synthetized and inserted into the pGADT7 plasmid (Genescript; Supplementary Figure 4). The plasmids pGADT7-PbRpt3-HA mutated 305-RKAEA-309 and pGADT7-PbRpt3-HA substituted for alanines in positions K188, D241, N288 and Q356 both optimized for *Xenopus laevis* were purchased from Azenta (Supplementary Figure 4). For the other mutations on pGAD-T7-PbRpt3, site-directed mutations were performed using the NEB Q5 Hot Start High-Fidelity DNA Polymerase (New England Biolabs, M0493) with primers generated via the NEBaseChanger tool (https://nebasechanger.neb.com/) and listed in Supplementary Table 2. This strategy was used for mutating 200-KVTF-203 in 200-KATA-203 on pGADT7-PbRpt3 (primers KA3 and KA4), and to obtain the double mutant, the mutation 305-RKAEA-309 was performed on pGADT7-PbRpt3-KATA mutated (primers RK3 and RK4). The PCR reactions were performed as recommended by the manufacturer (New England Biolabs) with optimized annealing temperatures as indicated in Supplementary Table 2. After analysis, 1µl of each PCR product was treated with KLD (kinase, ligase, DpnI) enzyme mix, and 5 µl of the treated mix were used to transform competent bacteria (Takara, Stellar™ Competent Cells, cat # 636763). Mutants were checked by sequencing and all primers used in this study are indicated in Supplementary Table 2.

### Recombinant protein expression

The coding regions of PbRpt3-45 kDa (a.a. 1-395) and PbRpt3-30 kDa (a.a.131-395) were obtained by PCR with the primers P1-P2 and P2-P4 respectively (Supplementary Table 2) and cloned into pETDuet-1 (Novagen) using the In-Fusion HD Cloning system (Clontech). All recombinant protein constructs were verified by sequencing. Recombinant His-tagged PbRpt3 expression was carried out in Artic Express BL21 Star™ (DE3) Chemically Competent *E. coli* cells (Life Technologies) in the presence of 0.5 mM IPTG at 10°C for 24 hours. After centrifuging at 3900 rpm for 30min at 4°C, cells were frozen at - 80°C overnight. Cells were resuspended in non-denaturing buffer at 4°C (10 mM Tris, 500 mM NaCl, MgCl2 1mM, 3U of DNase I, and protease inhibitor cocktail 1 X from Roche, ref 4693116001, pH 7.9) followed by sonication and ultracentrifugation at 13000 rpm for 45 min at 4°C. Pellets were resuspended in denaturing buffer (10 mM Tris, 500 mM NaCl, 6 M guanidine, 20 mM imidazole, MgCl2 1 mM and 1X protease inhibitor, pH 7.9) and sonicated again. After ultracentrifugation at 18000 rpm 45 min at 4°C, supernatant was incubated overnight at 4°C with Ni-NTA agarose beads (Qiagen) used to purify the recombinant proteins as described [40]. Beads were centrifuged at 3900 rpm for 20 min at 4°C and washed 4 times in denaturing buffer. Proteins were eluted with 12 ml of elution buffer (20 mM Tris, 500 mM NaCl, 6 M guanidine, 600 mM imidazole, 1 mM MgCl2 and 1X Roche protease inhibitor) and exchanged into #1 to #5 dialysis buffers in order to remove the imidazole and the Guanidine. Exchange buffers were prepared with 10% glycerol deionized water, 500 mM NaCl, 10 mM TrisHCl, 1mM MgCl2, pH 7.9. The exchange buffers #1, #2 and #3 contain additionally Guanidine at a concentration of 6M, 4M and 2M respectively, and exchange buffers #4 to #5 are identical and Guanidine free. After exchange in buffer #4, proteins were ultracentrifuged at 13000 rpm for 10 min at 4°C and concentrated to 1.5 ml using centrifugal filters (Merck, Amicon Ultra-4). They were then exchanged into buffer #5 overnight at 4°C and aliquoted and stored at -80°C. Recombinant proteins were quantified with Pierce™ BCA Protein Assay Kit (Life Technologies) and the purity of the purified proteins was checked by SDS-PAGE and western blot using anti-His antibody (1:1000, Qiagen) as primary antibody, HRP-labelled anti-mouse IgG (1:20000 dilution, Rockland Immunochemicals) as secondary antibody and Chemiluminescence detection (SuperSignal™ West Dura Extended Duration Substrate, Life Technologies).

### Measurement of binding of PbRpt3 to PP1c

PfPP1c (PF3D7_1414400) share a high level (99%) of identity of amino acid (301/304) with PbPP1c sequence (PBANKA_1028300), therefore we used the PfPP1c recombinant protein that we previously purified [32]. To assess binding of recombinant PbRpt3-30kDa and PbRpt3-45kDa proteins to PfPP1c, an ELISA-based assay was used as previously described [33]. Nunc-Immuno TM Microwell Maxisorp plates (Sigma-Aldrich) were coated overnight at 4°C with 25pmol of either PbRpt3-30kDa, PbRpt3-45kDa or PfI2 as positive control, diluted in PBS. After five washes with 0.1% PBS-Tween, the plates were blocked with 200 µl/well of PBS containing 0.5% gelatin for 1 h at room temperature. Plates were then incubated at 37 °C for 2 h with different quantities (0.125 - 4 pmol) of biotinylated PfPP1c labeled with biotin-NHS (Calbiochem) in PBS-Tween 0.1%. The binding detection was performed using Streptavidin-HRP (1:150000 in 0.1% PBS Tween) and trimethylbenzidine substrate (Uptima) (100 µl/well). The reaction was stopped using 2 N HCl. An ELISA plate reader (Multiskan FC, Thermo Fisher Scientific) was used to measure the optical density at 450 nm and 570 nm. The difference between these two OD measures was used for analysis. BSA was used as control. The statistical significance was calculated with the Mann-Whitney U test for nonparametric data. p values < 0.05 were considered significant.

### Assays for effects of PbRpt3 on PfPP1 activity

To investigate the role of PbRpt3 on PfPP1c activity, p-nitrophenyl phosphate (pNPP) was used in an assay as a substrate as described previously [32]. Briefly, different quantities of PbRpt3-30-kDa or PbRpt3-45kDa (37.5 to 600 pmol) were preincubated for 30 min at 37 °C with 1 µg of PfPP1c. After adding the substrate, the plate was incubated 1h at 37 °C and the variation of phosphatase activity of PfPP1c in presence of PbRpt3 was measured by optical density at 405 nm. Results are presented as mean of increase or decrease of PP1 phosphatase activity in comparison with PfPP1c activity without PbRpt3.

### Analysis of *Xenopus* oocytes GVBD and protein immunoprecipitation

The five plasmids (pGADT7-PbRpt3, pGADT7-PbRpt3-KATA, pGADT7-PbRpt3-RKAEA, pGADT7-PbRpt3-KATA/RKAEA, and pGADT7-PbRpt3 ATP-binding sites mutant), were linearized with XhoI, and used as templates with the T7 mMessage mMachine kit (Ambion) to synthesize *in vitro* capped mRNA (cRNA) encoding full size 45 kDa native or mutated protein PbRpt3. *Xenopus* oocytes were micro-injected with 80 nl of water used as control, or with cRNA coding for PbRpt3 native / mutants (60 ng in 80nl). We also injected PfI2 cRNA as a positive control (60 ng in 80nl) [34]. Microinjections were followed by progesterone extracellular treatment when indicated (10 μM, 15 hours after micro-injection) and for the experiment Figure 3A. After 15 h the GVBD (germinal vesicle breakdown) was detected by the appearance of a white spot at the center of the animal pole [54]. Experiments were performed using 10 oocytes and repeated on two to three animals. For immunoprecipitation experiments, 20 oocytes were lysed in 200 µl of buffer PY (50 mM Hepes pH 7.4, 500 mM NaCl, 0.05% SDS, 5 mM MgCl2, 10 µg/ml aprotinin, 10 µg/ml soybean trypsin inhibitor, 1 mg/ml bovine serum albumin, 10 µg/ml leupeptin, 1 mM PMSF, 10 µg/ml benzamidine, 1 mM sodium vanadate) and centrifuged at 4°C for 15 min at 12 000 g. Supernatants were incubated with anti-XePP1 antibodies (Santa-Cruz Biotechnology, 1:200) in the presence of protein A-sepharose beads (20 µl of 50% bead solution, Sigma) for 1 hour at 4°C. After centrifugation, beads were washed 3 times with buffer PY, and immune complexes are eluted with 25 µl of Laemmli buffer 2X (Biorad) heated at 90°C for 5 min. For electrophoresis, 15 µl of proteins complex solution were separated by 4–20% SDS-PAGE gels (mini protean TGX, BioRad), blotted onto nitrocellulose, and probed with primary antibody anti-XePP1 (Santa-Cruz Biotechnology, 1:1500) or anti-HA (1:1500, Invitrogen) followed by anti-mouse or anti-rabbit secondary antibodies (1:10000, eBiosciences, Trueblot). Chemiluminescence detection was performed with ECL Select detection system (Amersham) on hyperfilm (MP, Amersham).

### Isolation of parasites stages

Parasite stages were isolated as previously described [55–57]. Trophozoite stage parasites were obtained from blood collected by cardiac puncture from euthanized mice at a parasitaemia < 5%. For immunofluorescence experiments, mature schizonts were obtained from blood cultured (20 hours of culture) in a RPMI 1640 medium containing 25 mM HEPES, 0.4% Albumax, 0.2 mM hypoxanthine and 20 µg/ml gentamycin. For immunoprecipitation experiment, a mixed schizonts population (early, middle and late schizonts) was obtained after 15 hours culture in the same medium. This mixed population was purified on a 60% Nycodenz gradient (27.6% w/v Nycodenz in 5 mM Tris-HCl pH 7.20, 3 mM KCl, 0.3 mM EDTA) and centrifuged for 20 min à 450 g. For selection of gametocytes, 200 µl of phenylhydrazine (6 mg/ml) were injected (by IP) in mice 3 days prior to parasite infection. On day 3 post-infection and for 2 days, mice were treated with sulfadiazine (Sigma) 20 mg/l in drinking water, then, on day 5 post infection, blood collection was performed by cardiac puncture on euthanized mice. For all parasite stages, the purity of the parasite preparations was checked on Giemsa-stained smears by microscopic examination.

### Generation of transgenic parasites

The *Plasmodium berghei* (Pb)ANKA-GFP parasites were a kind gift from Dr. O. Silvie (Université Pierre et Marie Curie, Paris, France) and the pG230 parasites were a kind gift from Dr. N. Philip (The University of Edinburgh, Edinburgh, UK). A PbRpt3 KO line was generated by double homologous recombination of a NotI-linearized PlasmoGEM vector (PbGEM-022521, Wellcome Sanger Institute) transfected into PbANKA-GFP parasites. The C-terminal m-Cherry tagged PbRpt3 line was generated by single homologous recombination: a 1228-bp region of PbRpt3 starting 289 bp downstream from the start codon and lacking a stop codon was inserted into pL1886 vector (Genescript). A silent mutation was inserted in the coding sequence to generate an AvrII single cutter restriction site (Figure 5), in order to linearize the plasmid before transfection into PbANKA-GFP strain. All transfections were performed as previously described [45,58,59]. Nycodenz-enriched schizonts were electroporated with 10 µg linearized DNA and IV-injected into naive mice. Positive transfectants were selected with pyrimethamine in drinking water (70 mg/L, TCI). Four independent transfections were performed to generate KO lines, with a total number of 6 recipient mice. KO transgenic parasites were cloned by limiting dilution and genotyped using diagnostic PCR (Supplementary Table 2, Supplementary Figure 6). Primers QCR2 - GW2 and GW1 – GT were used to detect the integration of the dhfr resistance cassette at the PbRpt3 locus (5’- and 3’ side respectively), and primers QCR2 – P1 / QCR2 – P4 and/or QCR3 – P4 were used to detect the wild-type PbRpt3 gene. PbRpt3-mCherry parasites were enriched by cell-sorting on a FACSAria cell sorter (Beckton Dickinson) and the mCherry integration was confirmed by PCR using primers PF-877 (Supplementary Table 2). The correct size expression of PbRpt3-mCherry protein was confirmed by western blot analysis, with the parental strain used as control. Samples in Laemmli buffer were denatured at 100°C for 3 min and electrophoresed on a 4– 20% SDS polyacrylamide gel. Proteins were transferred onto nitrocellulose membranes (Amersham Biosciences). Membranes were probed with rabbit anti-RFP pAb (1:2000, MBL, PM005) as primary antibody, and anti-rabbit HRP (1:20000, Sigma-Aldrich, A0545) as secondary antibody. Chemiluminescence detection was performed as above.

### Immunofluorescence assays

Blood was collected from mice infected with *Plasmodium berghei* PbRpt3-mCherry parasites. *P. berghei* blood stages were fixed with 4% paraformaldehyde and 0.075% glutaraldehyde for 10 min at 4°C and centrifuged at 2000 rpm 2 min at room temperature. Sedimentation, permeabilization and saturation steps were performed as described [15]. Then a 1:500 dilution in PBS BSA 1% of Anti-RFP pAb (MBL, PM005) was applied for 1h at 37°C. Following PBS washing, coverslips were incubated with anti-Rabbit IgG Cross-Adsorbed, Alexa Fluor 594 (Life Technologies, A11012) diluted at 1:1000 in PBS BSA 1% and parasitic nucleus were stained with DAPI at a concentration of 1μg/ml in PBS BSA 1%, for 1h at 37°C. The coverslips were mounted in Mowiol (3µl), and confocal imaging was performed using a Zeiss LSM880 microscope. Subsequent image treatment was performed with ImageJ software.

### Immunoprecipitation

Purified parasites (mixed schizont population) of parental wild-type parasites used as control and PbRpt3-mCherry were suspended with 50 mM Tris-HCl, 0.5% Triton X100 and protease inhibitor cocktail (Roche), pH 8. A 10 freeze-thaw cycles and sonication 30’’ on/off cycles followed by a 5 hours centrifugation at 13000 rpm at 4° C allowed to obtain the soluble fractions. RFP-Trap1_A beads (Chromotek) were equilibrated with dilution buffer (20 mM Tris, 150 mM NaCl, 0.5% Triton X-100 and protease inhibitor cocktail (Roche), pH 7.5) and incubated overnight at 4°C with parasite soluble extracts on a rotating wheel. Beads were washed 10 times with dilution buffer and elution was performed in Laemmli buffer. After 3 min at 100°C, 8 samples were stored at -20°C for subsequent Mass-spectrometry experience (4 biological samples for Schizonts (15h culture) PbWT-GFP and 4 biological samples for Schizonts (15h culture) PbRpt3-mCherry). The presence of PbRpt3-mCherry in the IP was checked by western blot analysis as above.

### Sample preparation for mass spectrometry

S-Trap^TM^ micro spin column (Protifi, Hutington, USA) digestion was performed on immunoprecipitation eluates according to manufacturer’s instructions. Briefly, samples were supplemented with 20% SDS to a final concentration of 5%, reduced with 20mM TCEP (Tris(2-carboxyethyl) phosphine hydrochloride) and alkylated with 50mM CAA (chloracetamide) for 5min at 95°C. Aqueous phosphoric acid was then added to a final concentration of 2.5% following by the addition of S-Trap binding buffer (90% aqueous methanol, 100mM TEAB, pH7.1). Mixtures were then loaded on S-Trap columns. Five washes were performed for thorough SDS elimination. Samples were digested with 3µg of trypsin (Promega, Madison, WI, USA) at 47°C for 2 h. After elution, peptides were vacuum dried.

### NanoLC-MS/MS protein identification and quantification

The tryptic peptides were resuspended in 35 µL and a volume of 3µL was injected on a nanoElute (Bruker Daltonics, Germany) HPLC (high-performance liquid chromatography) system coupled to a timsTOF Pro (Bruker Daltonics, Germany) mass spectrometer. HPLC separation (Solvent A : 0.1% formic acid in water; Solvent B : 0.1% formic acid in acetonitrile) was carried out at 400nL/min using a packed emitter column (C18, 25 cm×75μm 1.6μm) (Ion Optics, Australia) using a 15min gradient elution (2 to 17% solvent B during 8min; 17 to 25% during 2min; 25% to 37% during 2min; 37% to 95% for 1min and finally 95% for 2min to wash the column). Mass-spectrometric data were acquired using the parallel accumulation serial fragmentation (PASEF) acquisition method in DIA (Data Independent Analysis) mode. The measurements were carried out over the m/z range from 475 to 1000 Th. The range of ion mobilities values from 0.85 to 1.30 V s/cm2 (1/k0). The total cycle time was set to 0.95s.

Data analysis was performed using DIA-NN software (version 1.8.1) [60]. A search against the UniProt/Swiss-Prot *Mus Musculus* database downloaded from Uniprot on 09/01/2023 (17574entries) and *Plasmodium berghei ANKA* database downloaded from PlasmoDB on 17/05/2023 (Release 63) was performed using library free workflow. For this purpose, “FASTA digest for library free search/library generation” and “Deep learning spectra, RTs and IMs prediction” options were checked for precursor ion generation. A maximum of 1 trypsin missed cleavages was allowed and the maximum variable modification was set to 2. Carbamidomethylation (Cys) was set as the fixed modification, whereas protein N-terminal methionine excision, methionine oxidation and N-terminal acetylation were set as variable modifications. The peptide length range was set to 7–30 amino acids, precursor charge range 2–4, precursor m/z range 300–1800, and fragment ion m/z range 300–1800. To search the parent mass and fragment ions, accuracy was set to 10 ppm manually. The false discovery rates (FDRs) at the protein and peptide level were set to 1%. Match between runs was allowed. For the quantification strategy, Robust LC (high precision) was used as advised in the software documentation, and normalization option was disabled whereas default settings were kept for the other algorithm parameters.

Statistical and bioinformatic analysis including volcano plot with Perseus were performed with Perseus software (version 1.6.15) [61] freely available at www.perseus-framework.org. All protein intensities were log2 transformed to perform statistics. For statistical comparison, we set two groups (WT, RPT3ch), each containing up to 4 biological replicates. We then filtered the data to keep only proteins with at least 4 out of 4 valid values in at least one group. Next, the data were imputed to fill missing data points by creating a Gaussian distribution of random numbers with a standard deviation of 33% relative to the standard deviation of the measured values and 2.5 standard deviation downshift of the mean to simulate the distribution of low signal values. We performed a t-test, FDR<0.01, S0=2.

## Author Contributions

Conceptualization, CP; Data curation, CL, CDW, AM, AF, IM, ICG, KC, CP; Formal analysis, CL, CDW, CP; Investigation, CL, CDW, AM, AF, IM, ICG, KC, CP; Methodology, CL, CDW, AM, AF, IM, ICG, KC, CP; Project administration, CP; Supervision, CL, JK, CP; Validation, CL, CDW, AF, ICG, KC, CP; Writing— original draft, CL, JK, CP; Writing—review and editing, CL, CDW, AM, AF, ICG, KC, JK, CP; All authors have read and agreed to the published version of the manuscript.

## Data Availability Statement

The datasets for this study can be found in PRIDE.

## Acknowledgments

The authors would like to thank Sophie Lecher, Elizabeth Pradel and Stéphanie Caby for technical assistance.

## References

1. Pickart, C.M.; Cohen, R.E. Proteasomes and Their Kin: Proteases in the Machine Age. Nat. Rev. Mol. Cell Biol. 2004, 5, 177–187, doi:10.1038/nrm1336.

2. Enenkel, C.; Kang, R.W.; Wilfling, F.; Ernst, O.P. Intracellular Localization of the Proteasome in Response to Stress Conditions. J. Biol. Chem. 2022, 298, 102083, doi:10.1016/j.jbc.2022.102083.

3. Wójcik, C.; DeMartino, G.N. Intracellular Localization of Proteasomes. Int. J. Biochem. Cell Biol. 2003, 35, 579–589, doi:10.1016/s1357-2725(02)00380-1.

4. Guo, X. Localized Proteasomal Degradation: From the Nucleus to Cell Periphery. Biomolecules 2022, 12, 229, doi:10.3390/biom12020229.

5. Sharon, M.; Taverner, T.; Ambroggio, X.I.; Deshaies, R.J.; Robinson, C.V. Structural Organization of the 19S Proteasome Lid: Insights from MS of Intact Complexes. PLoS Biol. 2006, 4, e267, doi:10.1371/journal.pbio.0040267.

6. Kunjappu, M.J.; Hochstrasser, M. Assembly of the 20S Proteasome. Biochim. Biophys. Acta 2014, 1843, 2–12, doi:10.1016/j.bbamcr.2013.03.008.

7. Scruggs, S.B.; Zong, N.C.; Wang, D.; Stefani, E.; Ping, P. Post-Translational Modification of Cardiac Proteasomes: Functional Delineation Enabled by Proteomics. Am. J. Physiol. Heart Circ. Physiol. 2012, 303, H9–18, doi:10.1152/ajpheart.00189.2012.

8. Guo, X.; Huang, X.; Chen, M.J. Reversible Phosphorylation of the 26S Proteasome. Protein Cell 2017, 8, 255–272, doi:10.1007/s13238-017-0382-x.

9. Zong, C.; Gomes, A.V.; Drews, O.; Li, X.; Young, G.W.; Berhane, B.; Qiao, X.; French, S.W.; Bardag-Gorce, F.; Ping, P. Regulation of Murine Cardiac 20S Proteasomes: Role of Associating Partners. Circ. Res. 2006, 99, 372–380, doi:10.1161/01.RES.0000237389.40000.02.

10. Zhang, F.; Hu, Y.; Huang, P.; Toleman, C.A.; Paterson, A.J.; Kudlow, J.E. Proteasome Function Is Regulated by Cyclic AMP-Dependent Protein Kinase through Phosphorylation of Rpt6. J. Biol. Chem. 2007, 282, 22460–22471, doi:10.1074/jbc.M702439200.

11. Kikuchi, J.; Iwafune, Y.; Akiyama, T.; Okayama, A.; Nakamura, H.; Arakawa, N.; Kimura, Y.; Hirano, H. Co- and Post-Translational Modifications of the 26S Proteasome in Yeast. Proteomics 2010, 10, 2769–2779, doi:10.1002/pmic.200900283.

12. Li, N.; Zhang, Z.; Zhang, W.; Wei, Q. Calcineurin B Subunit Interacts with Proteasome Subunit Alpha Type 7 and Represses Hypoxia-Inducible Factor-1α Activity via the Proteasome Pathway. Biochem. Biophys. Res. Commun. 2011, 405, 468–472, doi:10.1016/j.bbrc.2011.01.055.

13. Zhang, W.; Wei, Q. Calcineurin Stimulates the Expression of Inflammatory Factors in RAW 264.7 Cells by Interacting with Proteasome Subunit Alpha Type 6. Biochem. Biophys. Res. Commun. 2011, 407, 668–673, doi:10.1016/j.bbrc.2011.03.071.

14. Guo, X.; Engel, J.L.; Xiao, J.; Tagliabracci, V.S.; Wang, X.; Huang, L.; Dixon, J.E. UBLCP1 Is a 26S Proteasome Phosphatase That Regulates Nuclear Proteasome Activity. Proc. Natl. Acad. Sci. U. S. A. 2011, 108, 18649–18654, doi:10.1073/pnas.1113170108.

15. Hollin, T.; De Witte, C.; Fréville, A.; Guerrera, I.C.; Chhuon, C.; Saliou, J.-M.; Herbert, F.; Pierrot, C.; Khalife, J. Essential Role of GEXP15, a Specific Protein Phosphatase Type 1 Partner, in Plasmodium Berghei in Asexual Erythrocytic Proliferation and Transmission. PLoS Pathog. 2019, 15, e1007973, doi:10.1371/journal.ppat.1007973.

16. De Witte, C.; Aliouat, E.M.; Chhuon, C.; Guerrera, I.C.; Pierrot, C.; Khalife, J. Mapping PP1c and Its Inhibitor 2 Interactomes Reveals Conserved and Specific Networks in Asexual and Sexual Stages of Plasmodium. Int. J. Mol. Sci. 2022, 23, 1069, doi:10.3390/ijms23031069.

17. Krishnan, K.M.; Williamson, K.C. The Proteasome as a Target to Combat Malaria: Hits and Misses. Transl. Res. 2018, 198, 40–47, doi:10.1016/j.trsl.2018.04.007.

18. Dogovski, C.; Xie, S.C.; Burgio, G.; Bridgford, J.; Mok, S.; McCaw, J.M.; Chotivanich, K.; Kenny, S.; Gnädig, N.; Straimer, J.;, et al. Targeting the Cell Stress Response of Plasmodium Falciparum to Overcome Artemisinin Resistance. PLoS Biol. 2015, 13, e1002132, doi:10.1371/journal.pbio.1002132.

19. Bridgford, J.L.; Xie, S.C.; Cobbold, S.A.; Pasaje, C.F.A.; Herrmann, S.; Yang, T.; Gillett, D.L.; Dick, L.R.; Ralph, S.A.; Dogovski, C.;, et al. Artemisinin Kills Malaria Parasites by Damaging Proteins and Inhibiting the Proteasome. Nat. Commun. 2018, 9, 3801, doi:10.1038/s41467-018-06221-1.

20. Wang, L.; Delahunty, C.; Fritz-Wolf, K.; Rahlfs, S.; Helena Prieto, J.; Yates, J.R.; Becker, K. Characterization of the 26S Proteasome Network in Plasmodium Falciparum. Sci. Rep. 2015, 5, 17818, doi:10.1038/srep17818.

21. Muralidharan, V.; Oksman, A.; Iwamoto, M.; Wandless, T.J.; Goldberg, D.E. Asparagine Repeat Function in a Plasmodium Falciparum Protein Assessed via a Regulatable Fluorescent Affinity Tag. Proc. Natl. Acad. Sci. U. S. A. 2011, 108, 4411–4416, doi:10.1073/pnas.1018449108.

22. Li, H.; Bogyo, M.; da Fonseca, P.C.A. The Cryo-EM Structure of the Plasmodium Falciparum 20S Proteasome and Its Use in the Fight against Malaria. FEBS J. 2016, 283, 4238–4243, doi:10.1111/febs.13780.

23. Yokoyama, D.; Saito-Ito, A.; Asao, N.; Tanabe, K.; Yamamoto, M.; Matsumura, T. Modulation of the Growth OfPlasmodium Falciparum in Vitroby Protein Serine/Threonine Phosphatase Inhibitors. Biochem. Biophys. Res. Commun. 1998, 247, 18–23, doi:10.1006/bbrc.1998.8730.

24. Guttery, D.S.; Poulin, B.; Ramaprasad, A.; Wall, R.J.; Ferguson, D.J.P.; Brady, D.; Patzewitz, E.-M.; Whipple, S.; Straschil, U.; Wright, M.H.;, et al. Genome-Wide Functional Analysis of Plasmodium Protein Phosphatases Reveals Key Regulators of Parasite Development and Differentiation. Cell Host Microbe 2014, 16, 128–140, doi:10.1016/j.chom.2014.05.020.

25. Paul, A.S.; Miliu, A.; Paulo, J.A.; Goldberg, J.M.; Bonilla, A.M.; Berry, L.; Seveno, M.; Braun-Breton, C.; Kosber, A.L.; Elsworth, B.;, et al. Co-Option of Plasmodium Falciparum PP1 for Egress from Host Erythrocytes. Nat. Commun. 2020, 11, 3532, doi:10.1038/s41467-020-17306-1.

26. Bollen, M. Combinatorial Control of Protein Phosphatase-1. Trends Biochem. Sci. 2001, 26, 426– 431, doi:10.1016/s0968-0004(01)01836-9.

27. Ceulemans, H.; Bollen, M. Functional Diversity of Protein Phosphatase-1, a Cellular Economizer and Reset Button. Physiol. Rev. 2004, 84, 1–39, doi:10.1152/physrev.00013.2003.

28. Fardilha, M.; Esteves, S.L.C.; Korrodi-Gregório, L.; da Cruz e Silva, O. a. B.; da Cruz e Silva, F.F. The Physiological Relevance of Protein Phosphatase 1 and Its Interacting Proteins to Health and Disease. Curr. Med. Chem. 2010, 17, 3996–4017, doi:10.2174/092986710793205363.

29. Hendrickx, A.; Beullens, M.; Ceulemans, H.; Den Abt, T.; Van Eynde, A.; Nicolaescu, E.; Lesage, B.; Bollen, M. Docking Motif-Guided Mapping of the Interactome of Protein Phosphatase-1. Chem. Biol. 2009, 16, 365–371, doi:10.1016/j.chembiol.2009.02.012.

30. Bollen, M.; Peti, W.; Ragusa, M.J.; Beullens, M. The Extended PP1 Toolkit: Designed to Create Specificity. Trends Biochem. Sci. 2010, 35, 450–458, doi:10.1016/j.tibs.2010.03.002.

31. Choy, M.S.; Hieke, M.; Kumar, G.S.; Lewis, G.R.; Gonzalez-DeWhitt, K.R.; Kessler, R.P.; Stein, B.J.; Hessenberger, M.; Nairn, A.C.; Peti, W.;, et al. Understanding the Antagonism of Retinoblastoma Protein Dephosphorylation by PNUTS Provides Insights into the PP1 Regulatory Code. Proc. Natl. Acad. Sci. U. S. A. 2014, 111, 4097–4102, doi:10.1073/pnas.1317395111.

32. Daher, W.; Browaeys, E.; Pierrot, C.; Jouin, H.; Dive, D.; Meurice, E.; Dissous, C.; Capron, M.; Tomavo, S.; Doerig, C.;, et al. Regulation of Protein Phosphatase Type 1 and Cell Cycle Progression by PfLRR1, a Novel Leucine-Rich Repeat Protein of the Human Malaria Parasite Plasmodium Falciparum. Mol. Microbiol. 2006, 60, 578–590, doi:10.1111/j.1365-2958.2006.05119.x.

33. Fréville, A.; Landrieu, I.; García-Gimeno, M.A.; Vicogne, J.; Montbarbon, M.; Bertin, B.; Verger, A.; Kalamou, H.; Sanz, P.; Werkmeister, E.;, et al. Plasmodium Falciparum Inhibitor-3 Homolog Increases Protein Phosphatase Type 1 Activity and Is Essential for Parasitic Survival*. J. Biol. Chem. 2012, 287, 1306–1321, doi:10.1074/jbc.M111.276865.

34. Fréville, A.; Cailliau-Maggio, K.; Pierrot, C.; Tellier, G.; Kalamou, H.; Lafitte, S.; Martoriati, A.; Pierce, R.J.; Bodart, J.-F.; Khalife, J. Plasmodium Falciparum Encodes a Conserved Active Inhibitor 2 for Protein Phosphatase Type 1: Perspectives for Novel Anti-Plasmodial Therapy. BMC Biol. 2013, 11, 80, doi:10.1186/1741-7007-11-80.

35. Lenne, A.; De Witte, C.; Tellier, G.; Hollin, T.; Aliouat, E.M.; Martoriati, A.; Cailliau, K.; Saliou, J.-M.; Khalife, J.; Pierrot, C. Characterization of a Protein Phosphatase Type-1 and a Kinase Anchoring Protein in Plasmodium Falciparum. Front. Microbiol. 2018, 9, 2617, doi:10.3389/fmicb.2018.02617.

36. Yedidi, R.S.; Wendler, P.; Enenkel, C. AAA-ATPases in Protein Degradation. Front. Mol. Biosci. 2017, 4, 42, doi:10.3389/fmolb.2017.00042.

37. Inobe, T.; Genmei, R. N-Terminal Coiled-Coil Structure of ATPase Subunits of 26S Proteasome Is Crucial for Proteasome Function. PloS One 2015, 10, e0134056, doi:10.1371/journal.pone.0134056.

38. Dong, Y.; Zhang, S.; Wu, Z.; Li, X.; Wang, W.L.; Zhu, Y.; Stoilova-McPhie, S.; Lu, Y.; Finley, D.; Mao, Y. Cryo-EM Structures and Dynamics of Substrate-Engaged Human 26S Proteasome. Nature 2019, 565, 49–55, doi:10.1038/s41586-018-0736-4.

39. Khalife, J.; Fréville, A.; Gnangnon, B.; Pierrot, C. The Multifaceted Role of Protein Phosphatase 1 in Plasmodium. Trends Parasitol. 2021, 37, 154–164, doi:10.1016/j.pt.2020.09.003.

40. Fréville, A.; Cailliau-Maggio, K.; Pierrot, C.; Tellier, G.; Kalamou, H.; Lafitte, S.; Martoriati, A.; Pierce, R.J.; Bodart, J.-F.; Khalife, J. Plasmodium Falciparum Encodes a Conserved Active Inhibitor 2 for Protein Phosphatase Type 1: Perspectives for Novel Anti-Plasmodial Therapy. BMC Biol. 2013, 11, 80, doi:10.1186/1741-7007-11-80.

41. Tellier, G.; Lenne, A.; Cailliau-Maggio, K.; Cabezas-Cruz, A.; Valdés, J.J.; Martoriati, A.; Aliouat, E.M.; Gosset, P.; Delaire, B.; Fréville, A.;, et al. Identification of Plasmodium Falciparum Translation Initiation EIF2β Subunit: Direct Interaction with Protein Phosphatase Type 1. Front. Microbiol. 2016, 7, 777, doi:10.3389/fmicb.2016.00777.

42. Huchon, D.; Ozon, R.; Demaille, J.G. Protein Phosphatase-1 Is Involved in Xenopus Oocyte Maturation. Nature 1981, 294, 358–359, doi:10.1038/294358a0.

43. Egloff, M.P.; Johnson, D.F.; Moorhead, G.; Cohen, P.T.; Cohen, P.; Barford, D. Structural Basis for the Recognition of Regulatory Subunits by the Catalytic Subunit of Protein Phosphatase 1. EMBO J. 1997, 16, 1876–1887, doi:10.1093/emboj/16.8.1876.

44. Wakula, P.; Beullens, M.; Ceulemans, H.; Stalmans, W.; Bollen, M. Degeneracy and Function of the Ubiquitous RVXF Motif That Mediates Binding to Protein Phosphatase-1. J. Biol. Chem. 2003, 278, 18817–18823, doi:10.1074/jbc.M300175200.

45. Bushell, E.; Gomes, A.R.; Sanderson, T.; Anar, B.; Girling, G.; Herd, C.; Metcalf, T.; Modrzynska, K.; Schwach, F.; Martin, R.E.;, et al. Functional Profiling of a Plasmodium Genome Reveals an Abundance of Essential Genes. Cell 2017, 170, 260–272.e8, doi:10.1016/j.cell.2017.06.030.

46. Zhang, M.; Wang, C.; Otto, T.D.; Oberstaller, J.; Liao, X.; Adapa, S.R.; Udenze, K.; Bronner, I.F.; Casandra, D.; Mayho, M.;, et al. Uncovering the Essential Genes of the Human Malaria Parasite Plasmodium Falciparum by Saturation Mutagenesis. Science 2018, 360, eaap7847, doi:10.1126/science.aap7847.

47. Li, H.; O’Donoghue, A.J.; van der Linden, W.A.; Xie, S.C.; Yoo, E.; Foe, I.T.; Tilley, L.; Craik, C.S.; da Fonseca, P.C.A.; Bogyo, M. Structure- and Function-Based Design of Plasmodium-Selective Proteasome Inhibitors. Nature 2016, 530, 233–236, doi:10.1038/nature16936.

48. Otto, T.D.; Böhme, U.; Jackson, A.P.; Hunt, M.; Franke-Fayard, B.; Hoeijmakers, W.A.M.; Religa, A.A.; Robertson, L.; Sanders, M.; Ogun, S.A.;, et al. A Comprehensive Evaluation of Rodent Malaria Parasite Genomes and Gene Expression. BMC Biol. 2014, 12, 86, doi:10.1186/s12915-014-0086-0.

49. Laporte, D.; Salin, B.; Daignan-Fornier, B.; Sagot, I. Reversible Cytoplasmic Localization of the Proteasome in Quiescent Yeast Cells. J. Cell Biol. 2008, 181, 737–745, doi:10.1083/jcb.200711154.

50. Castaño, J.G.; Mahillo, E.; Arizti, P.; Arribas, J. Phosphorylation of C8 and C9 Subunits of the Multicatalytic Proteinase by Casein Kinase II and Identification of the C8 Phosphorylation Sites by Direct Mutagenesis. Biochemistry 1996, 35, 3782–3789, doi:10.1021/bi952540s.

51. Ebrahimzadeh, Z.; Mukherjee, A.; Crochetière, M.-È.; Sergerie, A.; Amiar, S.; Thompson, L.A.; Gagnon, D.; Gaumond, D.; Stahelin, R.V.; Dacks, J.B.;, et al. A Pan-Apicomplexan Phosphoinositide Binding Protein Acts in Malarial Microneme Exocytosis. EMBO Rep. 2019, 20, e47102, doi:10.15252/embr.201847102.

52. Maurya, R.; Tripathi, A.; Kumar, M.; Antil, N.; Yamaryo-Botté, Y.; Kumar, P.; Bansal, P.; Doerig, C.; Botté, C.Y.; Prasad, T.S.K.;, et al. PI4-Kinase and PfCDPK7 Signaling Regulate Phospholipid Biosynthesis in Plasmodium Falciparum. EMBO Rep. 2022, 23, e54022, doi:10.15252/embr.202154022.

53. Halgren, T.A. MMFF VI. MMFF94s Option for Energy Minimization Studies. J. Comput. Chem. 1999, 20, 720–729, doi:10.1002/(SICI)1096-987X(199905)20:7<720::AID-JCC7>3.0.CO;2-X.

54. Daher, W.; Oria, G.; Fauquenoy, S.; Cailliau, K.; Browaeys, E.; Tomavo, S.; Khalife, J. A Toxoplasma Gondii Leucine-Rich Repeat Protein Binds Phosphatase Type 1 Protein and Negatively Regulates Its Activity. Eukaryot. Cell 2007, 6, 1606–1617, doi:10.1128/EC.00260-07.

55. Beetsma, A.L.; van de Wiel, T.J.; Sauerwein, R.W.; Eling, W.M. Plasmodium Berghei ANKA: Purification of Large Numbers of Infectious Gametocytes. Exp. Parasitol. 1998, 88, 69–72, doi:10.1006/expr.1998.4203.

56. Rodríguez, M.C.; Margos, G.; Compton, H.; Ku, M.; Lanz, H.; Rodríguez, M.H.; Sinden, R.E. Plasmodium Berghei: Routine Production of Pure Gametocytes, Extracellular Gametes, Zygotes, and Ookinetes. Exp. Parasitol. 2002, 101, 73–76, doi:10.1016/s0014-4894(02)00035-8.

57. Fréville, A.; Gnangnon, B.; Tremp, A.Z.; De Witte, C.; Cailliau, K.; Martoriati, A.; Aliouat, E.M.; Fernandes, P.; Chhuon, C.; Silvie, O.;, et al. Plasmodium Berghei Leucine-Rich Repeat Protein 1 Downregulates Protein Phosphatase 1 Activity and Is Required for Efficient Oocyst Development. Open Biol. 2022, 12, 220015, doi:10.1098/rsob.220015.

58. Manzoni, G.; Briquet, S.; Risco-Castillo, V.; Gaultier, C.; Topçu, S.; Ivănescu, M.L.; Franetich, J.-F.; Hoareau-Coudert, B.; Mazier, D.; Silvie, O. A Rapid and Robust Selection Procedure for Generating Drug-Selectable Marker-Free Recombinant Malaria Parasites. Sci. Rep. 2014, 4, 4760, doi:10.1038/srep04760.

59. Janse, C.J.; Ramesar, J.; Waters, A.P. High-Efficiency Transfection and Drug Selection of Genetically Transformed Blood Stages of the Rodent Malaria Parasite Plasmodium Berghei. Nat. Protoc. 2006, 1, 346–356, doi:10.1038/nprot.2006.53.

60. Demichev, V.; Messner, C.B.; Vernardis, S.I.; Lilley, K.S.; Ralser, M. DIA-NN: Neural Networks and Interference Correction Enable Deep Proteome Coverage in High Throughput. Nat. Methods 2020, 17, 41–44, doi:10.1038/s41592-019-0638-x.

61. Tyanova, S.; Temu, T.; Sinitcyn, P.; Carlson, A.; Hein, M.Y.; Geiger, T.; Mann, M.; Cox, J. The Perseus Computational Platform for Comprehensive Analysis of (Prote)Omics Data. Nat. Methods 2016, 13, 731–740, doi:10.1038/nmeth.3901.

