## Supplementary Figures for "Characterization of the *Plasmodium berghei* regulatory AAA-ATPase subunit Rpt3 as an activator of Protein Phosphatase 1: direct and indirect evidence"

**SUPPLEMENTARY FIGURE 1.** Protein sequence of PbRpt3 (PBANKA_0715600)
**SUPPLEMENTARY FIGURE 2.** Sequence alignment of human PSMC4 with *Plasmodium berghei* PbRpt3.
**SUPPLEMENTARY FIGURE 3.** Analysis of PbRpt3 recombinant proteins.
**SUPPLEMENTARY FIGURE 4.** Recodonized sequence of PbRpt3 cDNA optimized for transcription in *Xenopus laevis*, and deduced amino acid sequence.
**SUPPLEMENTARY FIGURE 5.** PbRpt3 functional analysis using *Xenopus* oocytes model.
**SUPPLEMENTARY FIGURE 6.** PlasmoGEM construct (PbGEM-022521) and genotyping.
**SUPPLEMENTARY FIGURE 7.** STRING network visualization of PbRpt3-interacting proteins belonging to the 10 enriched GO terms.
**SUPPLEMENTARY FIGURE 8.** Ramachandran plot and statistics of the PbRpt3 model.

```
MRKMENVSKYLKEEDYYIKLKILKKQLDILNIQEEYIKEEHKNLKRELIRSKNEIKRIQS
VPLIIGQFLDIIDNNYGIVSSTAGSNYYVRILSTLNKEDLKPSVSVALHRHSHSIVNILP
SESDSSIQLLQVSERPNVKYTDLGGLDTQKQEMREAVELPLKSPELYEKIGIEPPMGILI
YGPPGTGKTMLVKAVANETKVTFIGVVGSEFVQKYLGEGPRMVRDVFRLARENSPSIIFI
DEVDAIATKRFDAQTGADREVQRILLELLNQMDGFDKSTNVKVIMATNRADTLDPALLRP
GRLDRKIEFPLPDRKQKRLIFQTIISKMNISSDVNIENFVVRTDKISAADIAAIAQESGM
QAIRKNRYIITASDFEQGYRTHVRKQLRDYEFYNI
```

**SUPPLEMENTARY FIGURE 1. Protein sequence of PbRpt3 (PBANKA_0715600)**. Two RVxF motifs are indicated in blue, the Walker A and Walker B motifs are highlighted in yellow.

```
                 Percent Identity  Matrix - created by Clustal2.1


                   1: HumanPSMC4_P43686_1      100.00   67.01
                   2: PBANKA_0715600_PbRpt3    67.01  100.00


CLUSTAL O(1.2.4) multiple sequence alignment


HumanPSMC4_P43686_1         MEEIGILVEKAQDEIPALSVSRPQTGLSFLGPEPEDLEDLYSRYKKLQQELEFLEVQEEY 60
PBANKA_0715600_PbRpt3       --------------------MRKMENV----SKYLKEEDYYIKLKILKKQLDILNIQEEY 36
                                                 *   .:     :  . ** * : * *:::*::*::****

HumanPSMC4_P43686_1         IKDEQKNLKKEFLHAQEEVKRIQSIPLVIGQFLEAVDQNTAIVGSTTGSNYYVRILSTID 120
PBANKA_0715600_PbRpt3       IKEEHKNLKRELIRSKNEIKRIQSVPLIIGQFLDIIDNNYGIVSSTAGSNYYVRILSTLN 96
                            **:*:****:*::::::*:*****:**:*****: :*:* .**.**:***********::

HumanPSMC4_P43686_1         RELLKPNASVALHKHSNALVDVLPPEADSSIMMLTSDQKPDVMYADIGGMDIQKQEVREA 180
PBANKA_0715600_PbRpt3       KEDLKPSVSVALHRHSHSIVNILPSESDSSIQLLQVSERPNVKYTDLGGLDTQKQEMREA 156
                            :* ***..*****:**:::*::** *:**** :*  .::*:* *:*:**:* ****:***

HumanPSMC4_P43686_1         VELPLTHFELYKQIGIDPPRGVLMYGPPGCGKTMLAKAVAHHTTAAFIRVVGSEFVQKYL 240
PBANKA_0715600_PbRpt3       VELPLKSPELYEKIGIEPPMGILIYGPPGTGKTMLVKAVANETKVTFIGVVGSEFVQKYL 216
                            *****.  ***::***:** *:*:***** *****.****:.*..:** ***********

HumanPSMC4_P43686_1         GEGPRMVRDVFRLAKENAPAIIFIDEIDAIATKRFDAQTGADREVQRILLELLNQMDGFD 300
PBANKA_0715600_PbRpt3       GEGPRMVRDVFRLARENSPSIIFIDEVDAIATKRFDAQTGADREVQRILLELLNQMDGFD 276
                            **************:**:*:******:*********************************

HumanPSMC4_P43686_1         QNVNVKVIMATNRADTLDPALLRPGRLDRKIEFPLPDRRQKRLIFSTITSKMNLSEEVDL 360
PBANKA_0715600_PbRpt3       KSTNVKVIMATNRADTLDPALLRPGRLDRKIEFPLPDRKQKRLIFQTIISKMNISSDVNI 336
                            :..***********************************:******.** ****:*.:*::

HumanPSMC4_P43686_1         EDYVARPDKISGADINSICQESGMLAVRENRYIVLAKDFEKAYKTVIKKDEQEHEFYK-  418
PBANKA_0715600_PbRpt3       ENFVVRTDKISAADIAAIAQESGMQAIRKNRYIITASDFEQGYRTHVRKQLRDYEFYNI  395
                            *::*.* ****.*** :*.***** *:*:****: *.***:.*:* ::*: :::***:
```

**SUPPLEMENTARY FIGURE 2. Sequence alignment of human PSMC4 (Uniprot reference P43686-1) with *Plasmodium berghei* PbRpt3 (PlasmoDB reference: PBANKA_0715600)**. Sequence alignment was performed using Clustal. An asterisk indicates positions which have a single, fully conserved residue. A double dot indicates conservation between groups of strongly similar properties (scoring > 0.5 in the Gonnet PAM 250 matrix), and a single dot indicates conservation between groups of weakly similar properties (scoring =< 0.5 and > 0 in the Gonnet PAM 250 matrix).

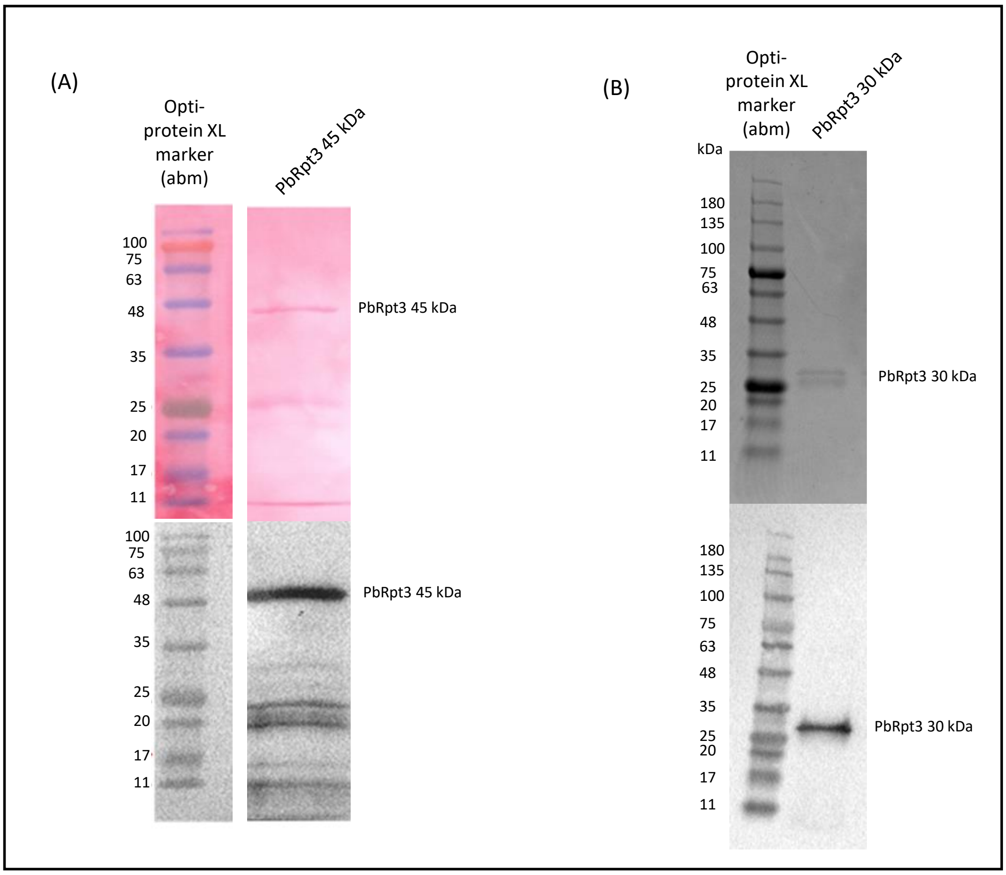


**SUPPLEMENTARY FIGURE 3. Analysis of PbRpt3-45kDa (A) and PbRpt3-30kDa (B) recombinant proteins.** The quality of PbRpt3-45kDa and PbRpt3-30kDa was checked by SDS-PAGE followed by rouge ponceau or coomassie blue coloration respectively (upper panel), and immunoblot analysis using an anti-His mAb (lower panel).

```
atg aga aaa atg gaa aac gtg agc aaa tac ttg aag gaa gag gac tac tac ata aaa ctg
 M   R   K   M   E   N   V   S   K   Y   L   K   E   E   D   Y   Y   I   K   L    20

aag ata ctg aag aaa caa ctt gac atc ctg aat att cag gaa gag tat atc aaa gaa gag
 K   I   L   K   K   Q   L   D   I   L   N   I   Q   E   E   Y   I   K   E   E    40

cac aaa aat ctg aag agg gag ctg atc aga tct aag aac gaa atc aag aga att cag agc
 H   K   N   L   K   R   E   L   I   R   S   K   N   E   I   K   R   I   Q   S    60

gtg cca ctg att atc gga cag ttc ctg gat atc atc gat aac aac tac gga atc gtg tct
 V   P   L   I   I   G   Q   F   L   D   I   I   D   N   N   Y   G   I   V   S    80

agc aca gcc gga tct aat tat tac gtg aga att ctg agt act ctg aac aaa gag gat ctg
 S   T   A   G   S   N   Y   Y   V   R   I   L   S   T   L   N   K   E   D   L    100

aag cct tcc gtg agt gtg gct ctg cat aga cac tct cat agc att gtg aat atc ctg cca
 K   P   S   V   S   V   A   L   H   R   H   S   H   S   I   V   N   I   L   P    120

tcc gag agt gat tcc agt att cag ctg ctg cag gtg tct gaa agg cct aac gtg aaa tat
 S   E   S   D   S   S   I   Q   L   L   Q   V   S   E   R   P   N   V   K   Y    140

aca gat ctg gga gga ctg gat act cag aag cag gaa atg aga gag gct gtg gaa ctg cca
 T   D   L   G   G   L   D   T   Q   K   Q   E   M   R   E   A   V   E   L   P    160

ctg aaa agc cct gag ctg tat gaa aag att gga atc gag cca cct atg gga att ctg atc
 L   K   S   P   E   L   Y   E   K   I   G   I   E   P   P   M   G   I   L   I    180

tac gga cca cct gga aca gga aaa act atg ctg gtg aag gca gtg gcc aat gaa aca aaa
 Y   G   P   P   G   T   G   K   T   M   L   V   K   A   V   A   N   E   T   K    200

gtg act ttt att gga gtg gtg gga tct gag ttc gtg cag aag tac ctg gga gaa gga cca
 V   T   F   I   G   V   V   G   S   E   F   V   Q   K   Y   L   G   E   G   P    220

aga atg gtt agg gat gtg ttt aga ctg gca agg gag aac tct cct agc att atc ttc att
 R   M   V   R   D   V   F   R   L   A   R   E   N   S   P   S   I   I   F   I    240

gat gaa gtg gat gct atc gca aca aaa agg ttt gat gcc cag act gga gct gat agg gag
 D   E   V   D   A   I   A   T   K   R   F   D   A   Q   T   G   A   D   R   E    260

gtg cag aga att ctg ctg gaa ctg ctg aac cag atg gat gga ttt gat aaa agc aca aat
 V   Q   R   I   L   L   E   L   L   N   Q   M   D   G   F   D   K   S   T   N    280

gtg aag gtg atc atg gca aca aac aga gcc gat act ctg gac cca gcc ctg ctg agg cca
 V   K   V   I   M   A   T   N   R   A   D   T   L   D   P   A   L   L   R   P    300

gga agg ctg gat aga aag att gag ttc cca ctg cct gat agg aaa cag aag aga ctg atc
 G   R   L   D   R   K   I   E   F   P   L   P   D   R   K   Q   K   R   L   I    320

ttc cag aca atc atc tcc aag atg aac atc tct agc gat gtg aac atc gaa aac ttc gtt
 F   Q   T   I   I   S   K   M   N   I   S   S   D   V   N   I   E   N   F   V    340

gtg aga act gat aag att tcc gct gca gat att gcc gct atc gct cag gag agt gga atg
 V   R   T   D   K   I   S   A   A   D   I   A   A   I   A   Q   E   S   G   M    360

cag gca atc agg aag aac aga tac atc atc act gca agt gat ttc gaa caa ggt tac agg
 Q   A   I   R   K   N   R   Y   I   I   T   A   S   D   F   E   Q   G   Y   R    380

aca cac gtt agg aag caa ctt aga gac tac gaa ttt tac aac att tga
 T   H   V   R   K   Q   L   R   D   Y   E   F   Y   N   I   -                    395
```

**SUPPLEMENTARY FIGURE 4. Recodonized sequence of PbRpt3 cDNA optimized for transcription in *Xenopus laevis*, and deduced amino acid sequence.** In blue : nucleotides coding for RVxF motifs : KVTF codons AAA**GTGACTTTT** will be substituted by KATA codons AAA**GCGACTGCT**. RKIEF codons AGAAAG**ATTGAGTTC** will be substituted by RKAEA codons AGAAAG**GCAGAGGCA** for the simple mutant, and by AGAAAG**GCTGAGGCC** for the double mutant. In yellow : nucleotides coding for the four residues K188, D241, N288 et Q356 which bind or stabilize the ATP have been substituted by GCA which codes for alanine.

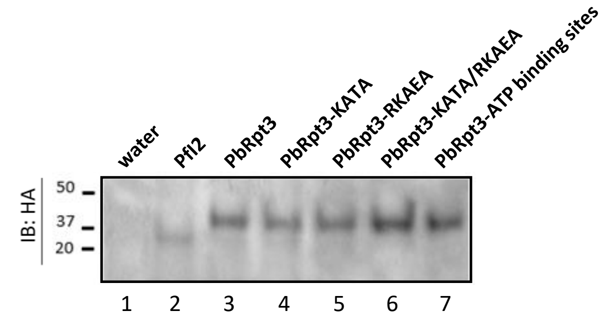


**SUPPLEMENTARY FIGURE 5. PbRpt3 functional analysis using *Xenopus* oocytes model**. PbRpt3 and mutated PbRpt3 proteins are expressed in *Xenopus* oocytes following the micro-injection of cRNA (60 ng in 80 nl). PfI2 is used as a positive control. Immunoblot analysis of extracts prepared from micro-injected oocytes with either water (lane 1) or cRNAs coding for PfI2 (lane 2), PbRpt3 (lane 3), PbRpt3-KATA (lane 4), PbRpt3-RKAEA (lane 5), PbRpt3-KATA/RKAEA (lane 6) or PbRpt3-ATP binding sites (lane 7) and followed by progesterone treatment (10 µM, 15 hours after micro-injection). The presence of proteins was detected using an anti-HA mAb.

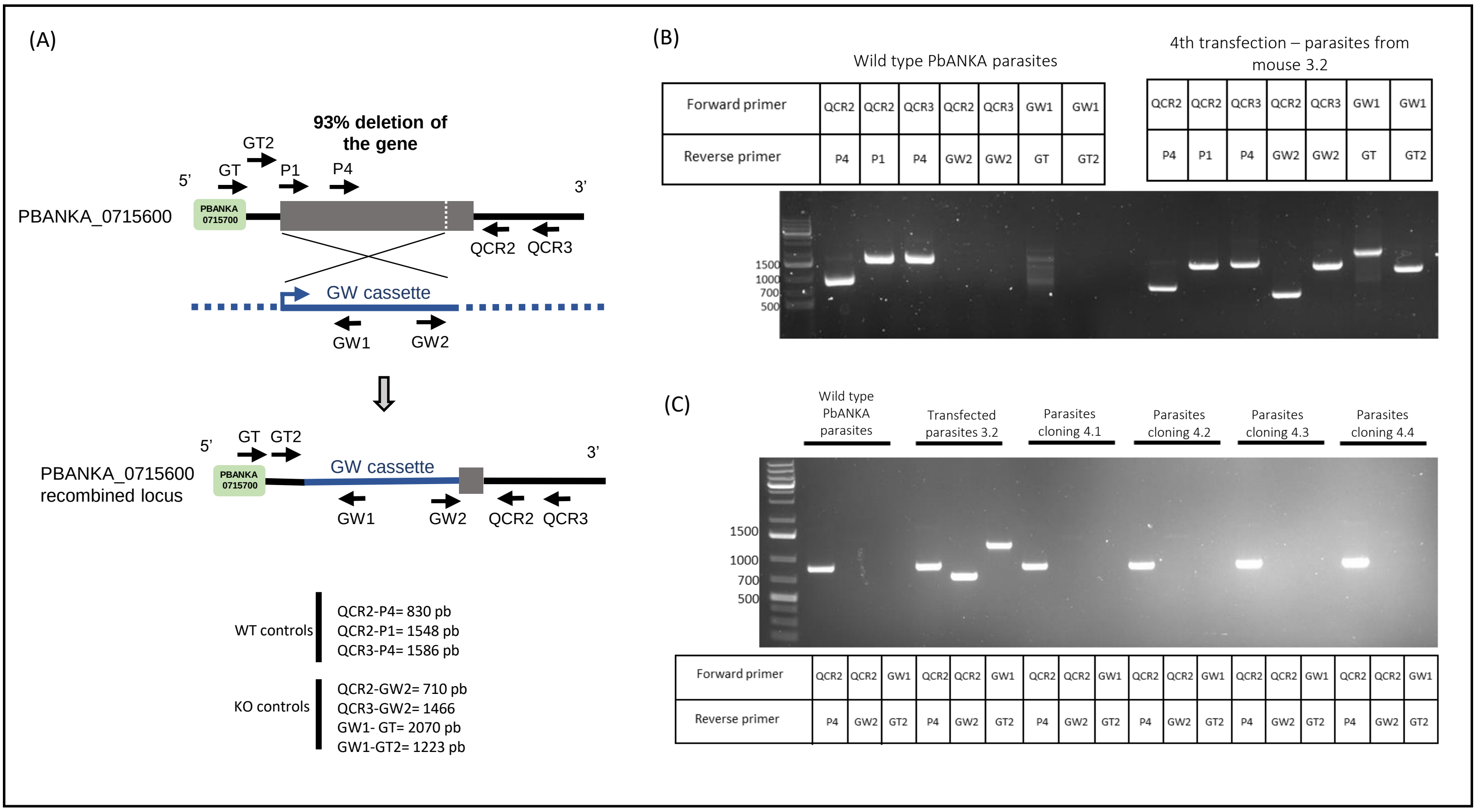


**SUPPLEMENTARY FIGURE 6. PlasmoGEM construct (PbGEM-022521) and genotyping. A.** The first panel shows PbRpt3 locus modification using the PbGEM-022521 construct provided by the Wellcome Sanger Institute. After linearization with NotI, the plasmid was transfected in PbANKA-GFP parasites, then injected to mice. The second panel indicates the sets of primers used and the expected fragments sizes : unmodified endogenous locus was detected using primers QCR2/P4, QCR2/P1 and/or QCR3/P4. The integration of the dhfr resistance cassette at the PbRpt3 locus was assessed using primers QCR2/GW2, QCR3/GW2, GW1/GT and/or GW1/GT2. **B.** PCR performed on wild-type (WT) parasites, or from parasites obtained in the 4th transfection (mouse 3.2). **C.** PCR performed on WT parasites, transfected parasites (mouse 3.2) or after cloning by limiting dilution in mice (mice 4.1 to 4.4).

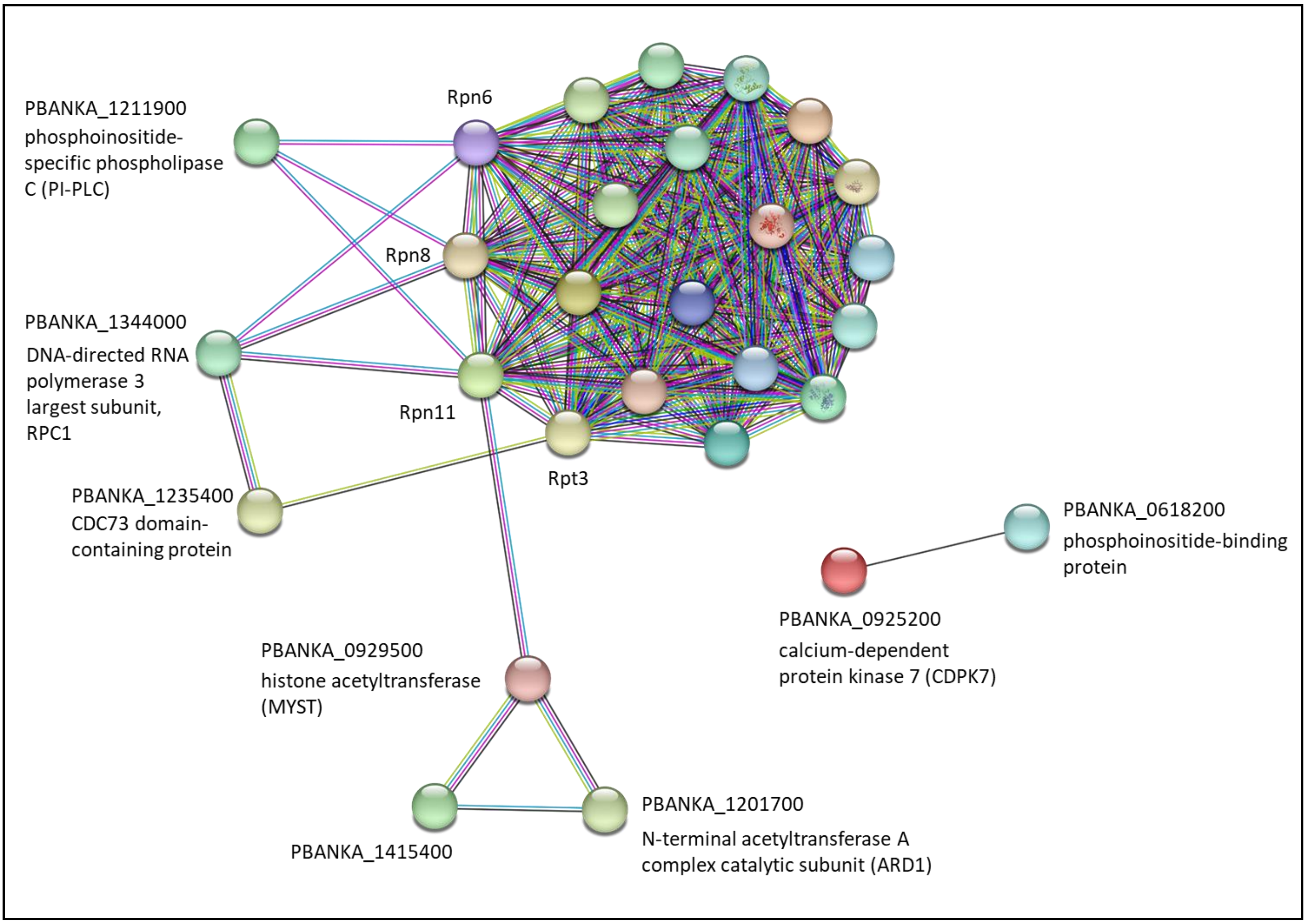


**SUPPLEMENTARY FIGURE 7. STRING network visualization of PbRpt3-interacting proteins belonging to the 10 enriched GO terms** (39 proteins, see Supplementary Table 3). With a protein–protein interaction enrichment *p*-value < 1.0e-16, the resulting network contained 203 functional associations among 28 proteins; the proteins with no associations to other proteins in the network were removed. For clarity, the gene Id and names of proteins belonging to the proteasome 19S were removed, except for Rpt3, Rpn6, Rpn8 and Rpn11 for which only the short names were kept. STRING analysis was performed using the 11.5 version of the software.

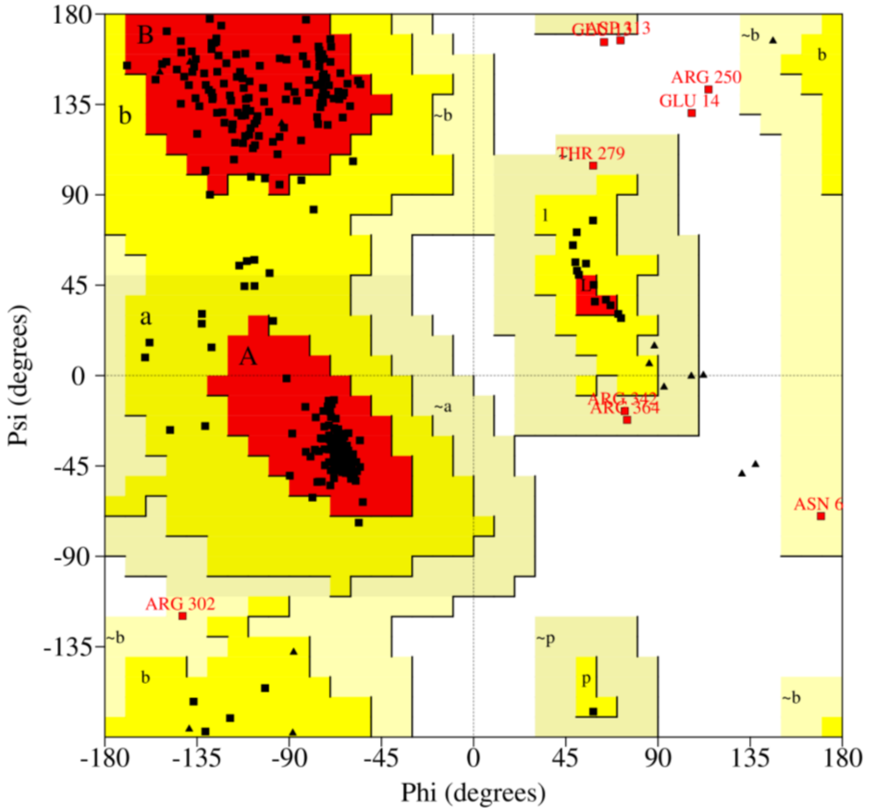


Plot statistics

| | | |
|---|---|---|
| Residues in most favoured regions [A,B,L] | 314 | 87.7% |
| Residues in additional allowed regions [a,b,l,p] | 35 | 9.8% |
| Residues in generously allowed regions [~a,~b,~l,~p] | 5 | 1.4% |
| Residues in disallowed regions | 4 | 1.1% |
|  | ---- | ------ |
| Number of non-glycine and non-proline residues | 358 | 100.0% |
| Number of end-residues (excl. Gly and Pro) | 2 | |
| Number of glycine residues (shown as triangles) | 19 | |
| Number of proline residues | 16 | |
|  | ---- |  |
| Total number of residues | 395 | |

**SUPPLEMENTARY FIGURE 8. Ramachandran plot and statistics of the PbRpt3** (PBANKA_0715600) **model**, based on the crystallographic structures of PSMC4  (PDB file 6MSB – chain D).
